# Concentration limits and localization of hydrogen peroxide in the extracellular space of solid tissues

**DOI:** 10.64898/2026.09.23.753516

**Authors:** Arthur Itacarambi, Marcos Gouveia, Rui D. M. Travasso, Armindo Salvador

## Abstract

H₂O₂ released to the extracellular space (ECS) regulates diverse physiological processes, yet its concentrations and spatial distribution in tissues remain poorly defined. This uncertainty hampers mechanistic understanding of redox signaling. Here, we used reaction–diffusion modeling to estimate extracellular H₂O₂ concentrations and transport ranges in various scenarios. Idealized analytical models were combined with numerical models incorporating localized NADPH oxidase (NOX) clusters, ECS microstructure, membrane permeability, and the thioredoxin- and GSH-dependent clearance systems. Using maximal neutrophil and NOX O_2_^•−^/H₂O₂ release rates, we obtained upper bounds for extracellular H₂O₂. Adjacent to isolated average-sized, fully active NOX2 clusters H_2_O_2_ peaked at ∼540 nM at adhesion cell–cell separations, and decreased radially over ∼50– 100 nm. At the receptor cell surface, peak concentration decreased inversely with intercellular separation, to <5 nM at 1 µm separation. Radial decrease here, for this wide separation, was over ∼2.5 µm. Even the former maximal extracellular concentrations induce just a minimal, highly localized oxidation of the intracellular Prdx, Trx and GSH pools. In turn, maximally activated neutrophils carry ∼2000 such NOX2 clusters, inducing 10’s of µM peak ECS H_2_O_2_ concentrations. These cause extensive Prdx and Trx oxidation near the exposed membranes. However, the GSH-dependent system still sustains a strong transmembrane gradient if the permeation barrier remains intact, and ECS H_2_O_2_ concentrations decay to sub-µM within a few µm of the source cell. Extracellular H_2_O_2_ concentrations scaled linearly with source flux in all the examined conditions. These results establish stringent constraints on autocrine, juxtacrine and next-cell paracrine H₂O₂ signaling.

## Introduction

Cellular release of superoxide (O_2_^•−^) and hydrogen peroxide (H_2_O_2_) to the extracellular space (ECS) regulates several important physiological and pathological processes [1–8]. However, the concentrations of these species in the ECS during signaling events remain a matter of speculation due to the lack of non-invasive probes with the required specificity, sensitivity and spatial resolution. Likewise, the spatial range of this signaling — whether autocrine, short-range paracrine, or long-distance volume transmission — remains under investigation [9–13]. Such information is critically important to understand the operation of the attending signal transduction mechanisms and for designing experimental protocols that more closely mimic the physiological environment [14,15].

Considering the shortcomings of currently available experimental methods, mathematical modeling presents an alternative to infer the concentration distribution and dynamics of extracellular reactive species from indirect experimental data (e.g. [13,16–24]). Through this approach, we have recently investigated the concentrations and transport ranges of O_2_^•−^ and H_2_O_2_ in the vasculature [13]. Of note, the results have shown that physiological H_2_O_2_ concentrations at the vasculature lumen adjacent to the vessel walls do not exceed the 10 nM range, much lower than the often claimed µM basal concentrations. The present work addresses the H_2_O_2_ concentrations and distribution in the ECS of non-vascular tissues. For that we use reaction-diffusion models that account for the five main determinants of these outcomes. Namely, (i) the release rates by cells, (ii) the spatial localization of release sites, (iii) the permeability of cell membranes, (iv) the ratio between the ECS volume and the surface of the cell plasma membranes in a tissue, and (v) the extent to which intracellular clearance outcompetes H_2_O_2_ efflux. Below we examine each of these factors in turn.

The highest O_2_^•−^/H_2_O_2_ release rates were recorded for phagocytic cells activated with phorbol myristate acetate (PMA). Converted to H_2_O_2_-equivalent amount ^1^, reported values are 0.13×10^6^ molecules s^−1^cell^−1^ for mouse J774A.1 macrophages at physiological (5%) O_2_ partial pressure [25], 20×10^6^ molecules s^−1^cell^−1^ for rat Kupffer cells [26], and (3.9 – 40)×10^6^ molecules s^−1^cell^−1^ for human neutrophils [27–29]. However, phagocytic cells treated with more-physiological stimulants and other cells typically show substantially lower rates: 0.34×10^6^ molecules s^−1^cell^−1^ for human neutrophils treated with opsonized zymosan [27], (0.060 – 0.19) × 10^6^ molecules s^−1^cell^−1^ for cultured bovine aortic endothelial cells at physiological O_2_ partial pressures [30,31], and values in the (0.0044 – 0.13) × 10^6^ molecules s^−1^cell^−1^ range for a variety of human, rat and mouse cells [32].

Because absent any strong oxidative stress the cytosolic antioxidant defenses clear virtually all the O_2_^•−^/H_2_O_2_ supplied to the cytosol (Supplementary Information Section 1 of ref. [33] for estimates), the rates above mainly reflect the release of these species to the ECS by NADPH oxidases (NOX) and dual oxidases (DUOX) at the cellular membrane [32]. Thus, the distribution of these enzymes over the cell membranes determines the extent of localization of O_2_^•−^/H_2_O_2_ release. The following evidence suggests that release usually occurs from localized spots at the cell surface. In vascular smooth muscle cells, NOX1 [34,35] and NOX5 [35] localize to caveolae, whereas NOX4 localizes to focal adhesions [34,35]. In human neutrophils, NOX2 localizes to lipid rafts at the cell membrane [36,37], where it is found in 0.03 – 0.1 µm^2^ clusters [38]. In thyrocytes, DUOXes localize to caveolae at apical microvilli [39], and at the membrane of bronchial epithelial cells DUOX1 also clusters at specific sites [40]. Whereas NOX4 [41,42] and DUOXes [43,44] release H_2_O_2_, the other NOXes release O_2_^•−^ [45], whose dismutation *via* the extracellular superoxide dismutase (SOD3) yields H_2_O_2_. In vascular tissues, SOD3 activity [46,47] can dismutate O_2_^•−^ within a fraction of a µm of the release sites. This may not be the case in tissues with lower SOD3 activity, but because SOD3 also localizes to lipid rafts / caveola [48–50] it is likely that a large fraction of the O_2_^•−^ produced therein is dismutated. However, O_2_^•−^ can also react very quickly with nitric oxide (*k* = 1.9×10^10^ M^−1^s^−1^ [51,52]), as well as permeate plasma membranes through anion channels [53–57]. Thus, even if O_2_^•−^ release is abundant and very localized, H_2_O_2_‘s yield and localization extent depend on these factors that remain poorly quantified.

Evidence for localized O_2_^•−^/H_2_O_2_ permeation is weaker. In human erythrocytes most H_2_O_2_ permeates passively across the lipid bilayer [58]. Because most reported plasma membrane permeabilities to H_2_O_2_ [59] are lower than human erythrocyte’s 16. µm s^−1^ [60], passive permeation may be the dominant mechanism in many other cell types as well. These observations imply that H_2_O_2_ permeation is, to a first approximation, relatively uniform across the cell surface. However, several lines of evidence support some degree of lateral heterogeneity. First, H_2_O_2_ permeability of lipid bilayers depends strongly on lipid composition [60], which varies across membrane regions such as lipid rafts and apical *vs*. basolateral. Secondly, multiple aquaporins (AQP0, 1, 3, 5, 6, 7, 8, 9, 11), collectively termed “peroxiporins”, facilitate H_2_O_2_ transport [7], and in some cells they significantly increase overall membrane permeability [61]. Their distribution within regions of otherwise low intrinsic H_2_O_2_ permeability may locally enhance H_2_O_2_ flux. Although AQP3 and AQP8 are enriched in caveolae [62–64], most cell membranes express additional peroxiporin variants [65] that are more uniformly distributed [66,67]. In contrast, O_2_^•−^ cannot significantly permeate lipid bilayers [53,54]. Instead, its transmembrane movement is mediated by anion channels [53–57]. While clustering of some of these channels has been documented in specific cell types [68,69], there is no evidence that such clustering is a general feature of plasma membranes. Taken together, these considerations indicate that although O_2_^•−^/H_2_O_2_ permeation is not strictly uniform over cell membranes, it is not usually restricted to discrete, highly localized sites.

Tissue architectures have narrow ECS domains where H_2_O_2_ could accumulate. Strikingly, widths of the ECS in the central nervous system are in the 30 – 60 nm range, with some pockets exceeding 100 nm [70,71]. The ECS of other tissues has been less scrutinized, but microscopy studies yielded the following results: at intercellular junctions the width of the ECS ranges from 10 nm, for tight junctions, to 25 nm for desmosomes; it increases to ∼35 nm between junctions [72]. However, intercellular junctions are crowded with protein, and most of the water in them is probably protein-bound. Because it is doubtful that H₂O₂ can diffuse appreciably through this space, we adopt 30 nm as the minimum cell–cell separation relevant for extracellular H₂O₂ accumulation and diffusion. Absent direct intercellular adhesion, extracellular matrix can separate cells by up to several µm [73].

The extent to which H_2_O_2_ clearance from the ECS is limited by membrane permeation or by the cytosolic clearance depends on whether the latter process outcompetes the H_2_O_2_ efflux (back-permeation). In the cytosol of human cells, the following three main enzymatic systems clear H_2_O_2_: the peroxiredoxin (Prdx) / thioredoxin (Trx) / thioredoxin reductase (TrxR) system, the glutathione peroxidase (GPx) / glutathione (GSH) / glutathione reductase (GSR) system, and catalase. The 2-Cys peroxiredoxins 1 and 2 (Prdx1/2) are both very abundant (adding up to >100 µM in most cells [74]) and very reactive with H_2_O_2_ (*k* ≍ 10^8^ M^−1^s^−1^ [75–78]), yielding estimated pseudo-first-order rate constants >10^4^ s^−1^. Thus, when fully reduced, these Prdx keep strong transmembrane (>200-fold) and cytosolic gradients ([79] and references therein). However, high H_2_O_2_ supply rates can substantially oxidize the Prdx pool. In their catalytic cycles these Prdx are first oxidized to the respective sulfenic acids (Prdx1/2-SOH), which then condense with another Cys in the active site, forming disulfides (Prdx1/2-SS). Thioredoxin 1 (Trx1-SH) reduces these disulfides and is thereby oxidized to its disulfide form (Trx1-SS). This is in turn reduced by NADPH under catalysis by thioredoxin reductase (TrxR). Computational modelling [33,74] and experimental evidence [80,81] indicate that the cytosolic TrxR activity is the main determinant of the maximal H_2_O_2_ supply rate that cells can sustain before the Prdx1/2-SH pools collapse. When the latter happens, GPx1 becomes the second line of defense. GPx1 catalyzes the reduction of H_2_O_2_ by GSH, but it is much less abundant in the cytosol (≍ 1 µM) than Prdx1/2, yielding pseudo-first-order rate constants for H_2_O_2_ reduction in the range 7 – 80 s^−1^ [74,82–86]. These can sustain transmembrane H_2_O_2_ gradients that, though more modest (≍ 5 – 20-fold), still make permeation the limiting factor in H_2_O_2_ clearance from the ECS. Oxidized glutathione (GSSG) is reduced by NADPH under catalysis by glutathione reductase and can also be actively excreted. Sustained very high oxidative loads may also deplete the GSH pool, and the last line of defense then becomes catalase. Catalase dismutates H_2_O_2_ and therefore does not directly depend on NADPH for clearing H_2_O_2_. However, in most human cells it is confined within peroxisomes which membrane limits H_2_O_2_ access [87]. This renders catalase unable to sustain a substantial H_2_O_2_ gradient across the plasma membrane, and under such strong stresses H_2_O_2_ clearance from the ECS may become limited by the effective catalase activity.

In the following sections, we introduce the numerical models and methods used. In the Results section, we first use analytical idealized reaction–diffusion models to gain generic insight on the interplay between tissue ECS microstructure, diffusion and membrane permeability. Leveraging on this insight we then analyze increasingly realistic numerical models that incorporate single or distributed NOX2 clusters, tissue ECS microstructure, membrane permeability, and explicit Prdx/Trx/TrxR and GPx/GSH/GSR networks in source and receptor cells. These models are used to map how cellular release rates, enzyme localization, tissue architecture and intracellular clearance together shape ECS H₂O₂ concentrations and signaling ranges. Finally, we discuss the implications for autocrine, juxtacrine and short-range paracrine redox signaling and oxidative stress in human tissues.

## Numerical models and methods

### Models

Below we use a series of idealized models with analytical steady state solutions (Models 1 - 3), as well as three more complex models (4 - 6) that can only be solved through numerical methods. Only the latter are described in the present section. The first of these models (**Model 4**) addresses the case where H_2_O_2_ emerges, with area-specific rate *ϕ_c_*, from the instantaneous dismutation of the O_2_^•−^ released by a circular NOX2 cluster of radius *R* at the surface of a cell. We considered the cylindrical geometry illustrated in Figure 1A (left) and assumed that (i) both membranes have a permeability constant *κ*, (ii) H_2_O_2_ clearance from the ECS is limited by the permeation barrier posed by the membranes, and (iii) there are no barriers to H_2_O_2_ diffusion within the ECS. The distribution of the H_2_O_2_ concentration over the radial (*r*) and axial (*x*) directions at steady state, [H_2_O_2_](*r*, *x*), is then given by:

**Figure 1.**
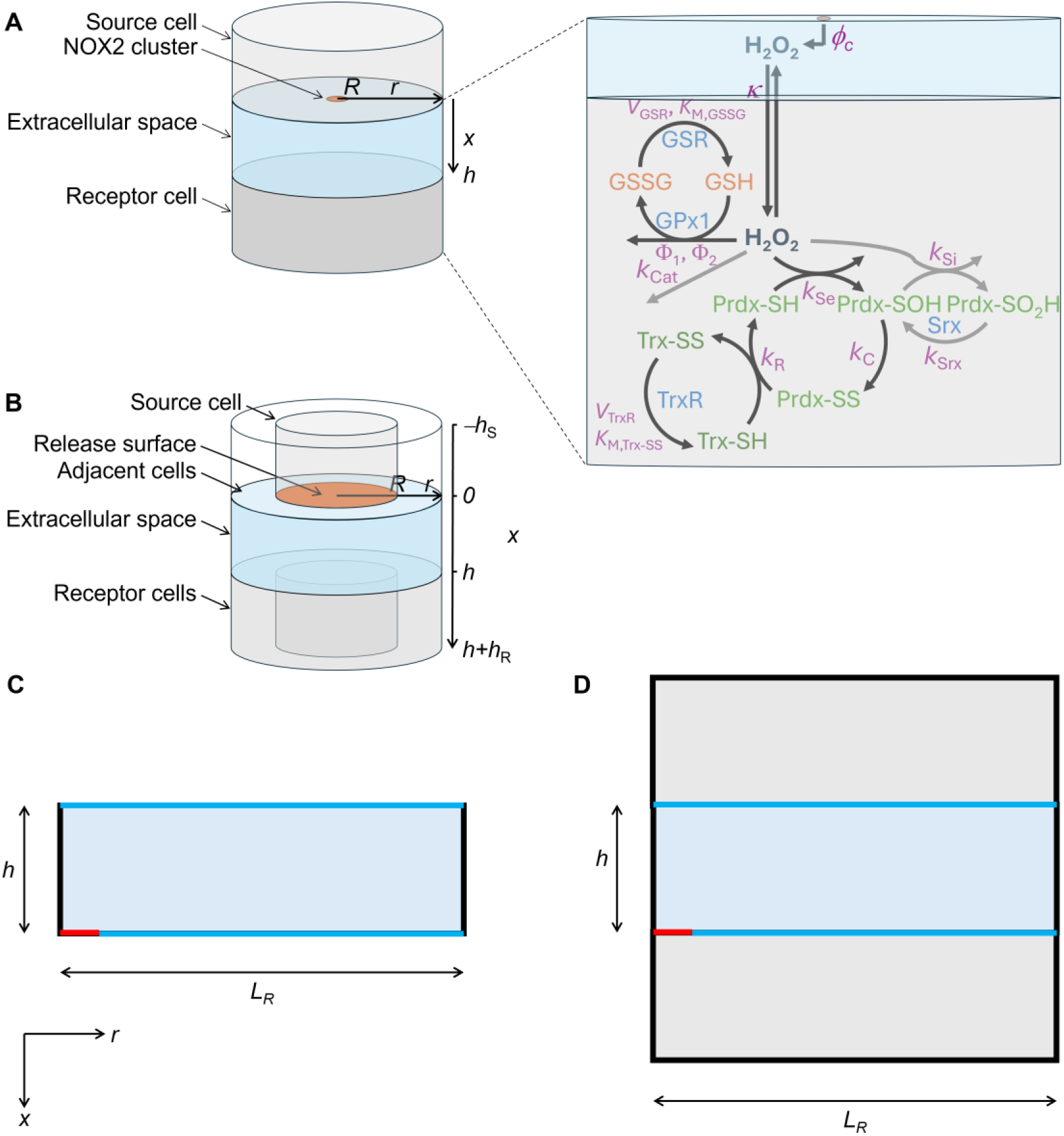
Models for H_2_O_2_ diffusion and metabolism. (**A**) We consider a cylindrical geometry as depicted (left). O_2_^•−^ is released to the ECS (cyan) from a circular NOX2 cluster (orange) at the membrane of a source cell (e.g., neutrophil) and instantaneously dismutated to H_2_O_2_. The latter permeates into both the source and receptor cells either irreversibly (**Model 4**), or reversibly with the back-flux in competition with metabolism in both these cells (**Model 5**). Metabolism involves both the Prdx/Trx/TrxR and the GSH/GPx/GSR systems (right). The enzymes catalyzing each process are shown in cyan, and the parameters of the rate laws in magenta. The reactions in gray are neglected in Model 5 but not in the next one. (**B**) Model considering that H_2_O_2_ is uniformly released throughout the membrane of the source cell facing the considered ECS segment (orange) and intracellular redox pools in both cylindrical central cells are confined within these (**Model 6**). H_2_O_2_ is considered to permeate both the membranes facing the ECS and those forming the radial wall of the central cells. (**C**) Diagram of simulated system for Model 4. The simulation is carried out in cylindrical coordinates in a two-dimensional revolution plane. (**D**) Diagram of simulated system for Model 5. In Models 5 and 6 new domains representing the cells are added. In these domains both the Prdx/Trx/TrxR and the GSH/GPx/GSR systems are simulated.

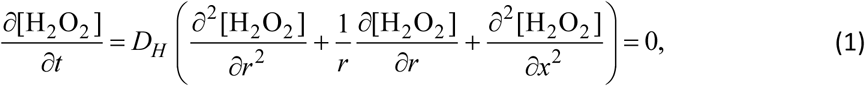

with the boundary conditions

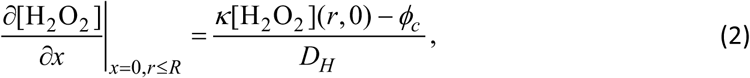

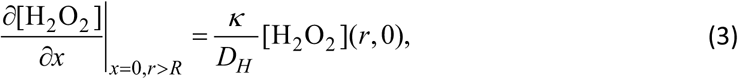

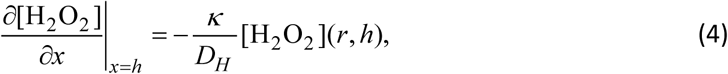

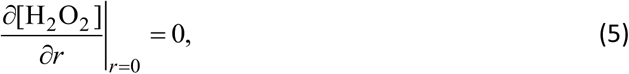

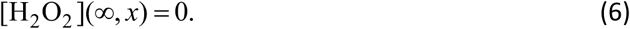

In the second model (**Model 5**, Figure 1A), assumption (ii) is relaxed, and the permeation rate depends on the H_2_O_2_ clearance by the cytosolic metabolism of both cells. For simplicity, we assumed that both cells have similar total concentrations of glutathione and of all the considered proteins: Prdx, Trx1, TrxR1, GPx1, GSR (Figure 1A, right). We used as reference the concentrations and activities in human hepatocytes, which are among the best studied human cells from solid tissues. We aggregated the two cytosolic 2-Cys Prdx (Prdx 1 and 2) into a single Prdx with intermediate properties, similarly to ref. [74]. We also neglected Prdx sulfinylation, which occurs very slowly, and a possible delay in recovering Prdx’s fully folded conformation after reduction by Trx1 [88]. For GPx1 we considered the Dalziel steady-state kinetics [89], as the rate constants of the individual steps in the catalytic cycle are unknown. For similar reasons, we considered the steady-state kinetics of glutathione reductase [90] and thioredoxin reductase [91,92]. Because these enzymes have *K*_M_ for NADPH (8.5 µM [93] and 6.0 µM [94], respectively) that are much lower than the NADPH concentration in these cells (3.0×10^2^ µM [95]) we considered them saturated with NADPH. We neglected the action of Cat, which in most cells contributes much less to cytosolic H_2_O_2_ clearance under physiological conditions.

For the cytosol we adopted the H₂O₂ diffusion coefficient (*D_H_* _,*cyt*_) measured in a hydrogel of comparable viscosity [96]; for the ECS we adopted the value in water [96], ∼5-fold higher. While the latter may be an overestimate, studies of H_2_O_2_ diffusion in the brain striatum of living rats support such a high diffusion coefficient [12]. For GSH and GSSG we adopted the diffusion coefficients estimated in ref. [97], based on those for similarly-sized molecules in cells [98]. Likewise, the adopted diffusion coefficients of the various Prdx and Trx1 redox forms are similar to those determined for similarly-sized molecules in the cytosol, as estimated in ref. [33], considering that Prdx remains decameric over its redox cycle [99].

The partial differential equations and boundary conditions describing the dynamics of this system are shown in Supplementary Information Section 1 (SI1) and the reference parameters values presented in Table 1.

**Table 1.** Symbol meanings and parameter values.

| **Symbol** | **Meaning** | **Value** | **Ref.** |
| --- | --- | --- | --- |
| $\phi$ | Max. area-specific H<sub>2</sub>O<sub>2</sub>-equiv. production rate of neutrophils | $1.8 \times 10^5 \text{ molec s}^{-1} \mu\text{m}^{-2}$ | See text |
| $\phi_c$ | Max. area-specific H<sub>2</sub>O<sub>2</sub> -equiv. production rate of a NOX2 cluster | $5.7 \times 10^6 \text{ molec s}^{-1} \mu\text{m}^{-2}$ | See text |
| $\kappa$ | Typical permeability constant of plasma membranes for H<sub>2</sub>O<sub>2</sub> | $10. \mu\text{m s}^{-1}$ | [74] |
| $h_S, h_R$ | Height of source and receptor cells | $5.6 \mu\text{m}, 11.2 \mu\text{m}$ | |
| $D_H$ | H<sub>2</sub>O<sub>2</sub> diffusion coefficient in the ECS | $1.83 \times 10^3 \mu\text{m}^2 \text{s}^{-1}$ | See text |
| $D_{H,cyt}$ | H<sub>2</sub>O<sub>2</sub> diffusion coef. in the cytosol | $3.7 \times 10^2 \mu\text{m}^2 \text{s}^{-1}$ | See text |
| $D_{\text{GSH}}$ | GSH diffusion coefficient | $1.8 \times 10^2 \mu\text{m}^2 \text{s}^{-1}$ | See text |
| $D_{\text{GSSG}}$ | GSSG diffusion coefficient | $80. \mu\text{m}^2 \text{s}^{-1}$ | See text |
| $D_{\text{P10}}$ | Prdx diffusion coefficient | $0.52 \mu\text{m}^2 \text{s}^{-1}$ | See text |
| $D_{\text{Trx}}$ | Trx1 diffusion coefficient | $35. \mu\text{m}^2 \text{s}^{-1}$ | See text |
| $k_{\text{Se}}$ | Prdx-SH sulfenylation rate constant | $100. \mu\text{M}^{-1} \text{s}^{-1}$ | [75–78] |
| $k_{\text{Si}}$ | Prdx-SH sulfinylation rate constant | $0.0019 \mu\text{M}^{-1} \text{s}^{-1}$ | [100–104] |
| $k_C$ | Prdx-SOH condensation rate ctt. | $8.7 \text{ s}^{-1}$ | [77,78,101,102] |
| $k_R$ | Prdx-SS reduction rate constant | $1.9 \mu\text{M}^{-1} \text{s}^{-1}$ | [105] |
| $k_{\text{Srx}}$ | Prdx-SO<sub>2</sub>H reduction rate ctt. | $2.9 \times 10^{-4} \text{ s}^{-1}$ | [74] |
| $V_{\text{TrxR}}$ | Maximal rate of Trx reductase | $50. \mu\text{M s}^{-1}$ | [74] |
| $K_{\text{M,Trx-SS}}$ | TrxR's Michaelis ctt. for Trx1-SS | $1.8 \mu\text{M}$ | [106] |
| [GPx1] | GSH peroxidase 1 concentration | $2.4 \mu\text{M}$ | [74] |
| $\Phi_1$ | Dalziel parameter of GPx1 for H<sub>2</sub>O<sub>2</sub> | $0.024 \mu\text{M s}$ | [107] |
| $\Phi_2$ | Dalziel parameter of GPx1 for GSH | $4.3 \mu\text{M s}$ | [107] |
| $V_{\text{GSR}}$ | Maximal rate of GSSG reductase | $195. \mu\text{M s}^{-1}$ | [108] |
| $K_{\text{M,GSSG}}$ | GSR's Michaelis ctt. for GSSG | $65. \mu\text{M}$ | [93] |
| [Prdx]<sub>T</sub> | Total conc. of Prdx monomers | $86. \mu\text{M}$ | [74] |
| [Trx]<sub>T</sub> | Total concentration of Trx1 | $63. \mu\text{M}$ | [74] |
| [GS]<sub>T</sub> | Total conc. of glutathionyl moieties | $8.0 \text{ mM}$ | [109] |

We also considered a model (**Model 6**) where H_2_O_2_ is uniformly released throughout the membrane of the source cell facing the considered ECS segment (Figure 1B). In this model, both the source and the receptor cell are cylindrical with the same radius (*R*). H_2_O_2_ permeates the membranes facing the shown ECS segment and those forming the cylinders’ radial walls. For simplicity, we consider that the latter membranes directly adjoin those of the radially surrounding cells without a ECS layer, and thus these together constitute a single permeation barrier with half the permeability constant of a single membrane. The intracellular redox pools cannot permeate any of the membranes. The cells surrounding the central ones metabolize H_2_O_2_ in the same way as these. For simplicity, we consider those cells unbounded in the radial direction for *r*>*R*. We set the height (*h*_S_) of the source cell and radially surrounding space to 5.6 µm, to match the volume of a neutrophil, and the height (*h*_R_) of the receptor cell as twice that value. The equations for this model are the same as for the previous one, except for the boundary conditions defining the radial limits of the source and receptor cells. The values of all the parameters other than *R* and *ϕ* are also the same for both models. In Model 6 the latter parameter represents the mean area-specific H_2_O_2_ release rate of the source cell, while in Models 4 and 5 we considered instead the H_2_O_2_ release rate of a NOX2 cluster, *ϕ_c_*. Model 6 accounts for Prdx sulfinylation and treats Prdx-SO_2_H reduction as a first-order process.

### Methods

The simulations were carried out in cylindrical coordinates. For Model 4 we consider a two-dimensional revolution plane with sufficiently large radial length *L_R_*, so that the calculated concentrations are approximately equal to zero for *r* = *L_R_* (in this setting we have verified that it was sufficient to set *L_R_* = 21 µm) and with variable axial length *h* (depending on the simulation). This domain is discretized considering the lattice spaces Δ*r* = Δ*x* = 5 nm. In Figure 1C we represent a diagram of this simulation domain where the H_2_O_2_ permeability condition (3) is imposed on the blue boundaries and the H_2_O_2_ production condition (2) is imposed on the red boundary. The black boundary at *r* = 0 has a zero-flux boundary condition (5), and the boundary at *r* = *L_R_* has a fixed value boundary condition (6), though the simulation results are the same for zero flux at that boundary. The system of equations was solved in MatLab in parallel with 12 processors using the function *ode45*.

For Models 5 and 6 we add two new domains representing the source and receptor cells (Figure 1D). In these domains we consider both the Prdx/Trx/TrxR and the GSH/GPx/GSR systems. For Model 5 we use *L_R_* = 12.5 µm with Δ*r* = 5 nm. In the ECS domain we set Δ*x* = 5 nm, while we set Δ*x* = 10 nm in the two cellular domains. For Model 6, each cellular domain is divided in two (with a zero-flux boundary condition), according to Figure 1B. The larger H_2_O_2_ production region leads to shallower concentration gradients and permits larger spatial increments. Therefore, we consider *L_R_* = 102 µm with Δ*r* = Δ*x* = 20 nm.

The simulations of Models 5 and 6 are too large to be run in MatLab, and we need to use an iterative solver. Here we use CVODE [110] from the Sundials (SUite of Nonlinear and DIfferential/ALgebraic equation Solvers) library [111]. CVODE iteratively obtains the solution of a very large system of equations using the generalized minimal residual method [112] coupled with an implicit backward differentiation formula to tackle stiff equations. Here the code was written in FORTRAN with OpenMP to run in parallel in 32 processors.

## Results

We begin with the simplest scenario (**Model 1**), by considering that H_2_O_2_ is uniformly released at an area-specific rate *ϕ* from one of the infinite parallel plane plasma membranes from two adjacent cells separated by a distance *h*. Assuming that H_2_O_2_ is only cleared from the ECS through permeation-limited absorption by the cells, and that both membranes have permeability constant *κ*, at steady state the rates of release and absorption must balance. We use this fact to estimate the H_2_O_2_ concentration. That is:

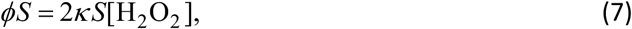

where *S* stands for the surface area of each membrane. Therefore, the steady state concentration will be simply:

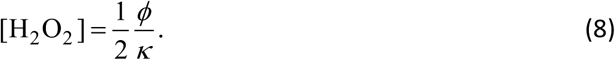

To obtain an upper bound we will consider the highest production rate recorded for a cell. Namely, we consider the H_2_O_2_-equivalent release of *P*=40×10^6^ molecules/s/cell determined for human neutrophils [29]. For a typical 4.2 µm radius neutrophil [113], one obtains:

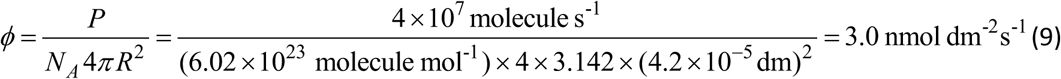

Considering *κ* = 10 µm s^−1^, as is typical for human cell lines [74] yields [H_2_O_2_] = (3.0 ×10^−9^ mol dm^-2^s^-1^) / (2 ×1.0 ×10^−4^ dm s^-1^) =15. μM.

Importantly, this value substantially overestimates the mean H_2_O_2_ concentrations that can be reached in the ECS for the following four reasons. First, the considered area-specific release rate represents a maximal rate achieved under artificial stimulation of the most productive human cells and is unlikely to be reached under most physiological conditions. Second, part of the released O_2_^•−^ might react with ^•^NO, be absorbed by the cells or diffuse away from the production zone, instead of being all dismutated in the ECS as assumed above. Third, in the physiological environment, conditions that propitiate abundant O_2_^•−^/H_2_O_2_ release by neutrophils also cause rapid release of myeloperoxidase [1,114], which catalyzes the consumption of a substantial fraction of the released H_2_O_2_ [29]. Fourth, and not least, we consider production over an infinite plane membrane, rather than over the surface of a small cell, and neglect leakage by diffusion over the ECS network extending over the tissue. We will examine the consequences of this fourth point below.

First we assess the amplitude of the concentration gradient that is expected in the present geometry. This gradient develops because H_2_O_2_ is released from just one of the membranes and absorbed by both membranes. In this situation, the H_2_O_2_ concentration at steady state ([H_2_O_2_](*x*)) and the gradient amplitude can be computed through the following diffusion model:

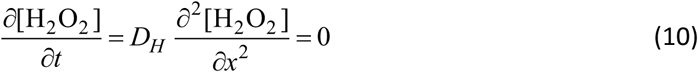

with boundary conditions

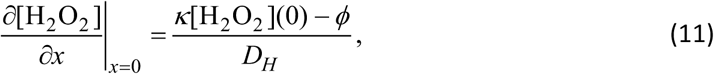

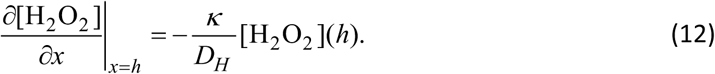

The steady state solution is:

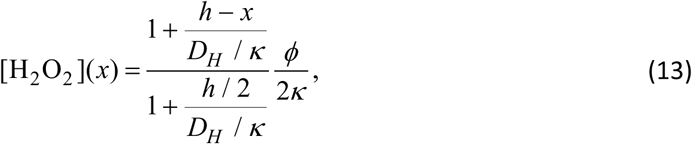

yielding the following expression for relative amplitude:

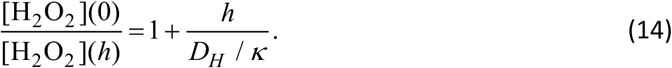

Considering *κ* = 10 µm s^−1^ as above and *D_H_* = 1830 µm^2^ s^−1^ [96] yields *D_H_* / *κ* = 183 μm, meaning that the gradient amplitude is negligible for physiologically relevant cell separations such as below 1 µm.

To assess the importance of geometry let us now consider that the H_2_O_2_ is released, instead, from a spherical cell of radius *R* surrounded by a spherical shell of cells separated from it by ECS of thickness *h* (**Model 2**). As before, we consider that the membranes of both the inner and the outer cells have permeability constant *κ*. The reaction-diffusion model for this system, expressed in spherical coordinates as function of the distance (*r*) to the center of the cell, is:

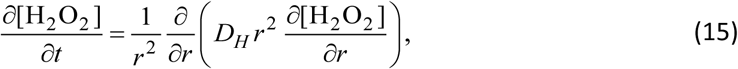

with boundary conditions

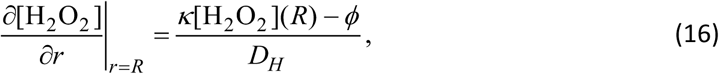

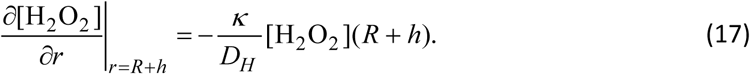

The steady state solution is:

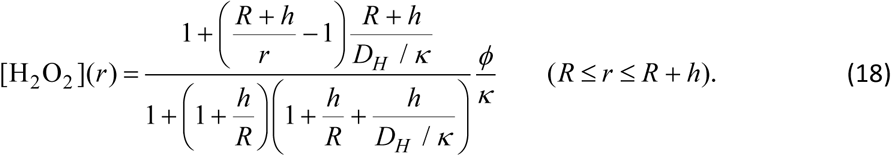

The estimated steady state concentrations obtained through this equation are slightly lower than those obtained for the planar geometry, especially for the higher separation between cells. Thus, considering *R*= 4.2 µm as the radius of a typical human neutrophil [113] and the same values of *D*_H_, *κ* and *ϕ* as above, for a separation *h*= 1 µm the concentration at the surface of the inner cell becomes 12. µM.

The relative amplitude of the gradient in this geometry is:

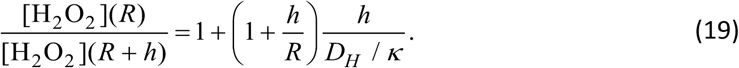

This equation reveals a quadratic dependence on the separation between cells, instead of a linear dependence found for a planar geometry. However, the gradient amplitude, though slightly higher, remains negligible for physiologically relevant cell separations: a mere 0.67% decrease for a 1 µm separation between the inner and the outer cells.

To assess the effect of H_2_O_2_ leakage over the ECS network surrounding the source cell we consider again a spherical cell of radius *R* uniformly releasing H_2_O_2_ from its surface at an area-specific rate *ϕ*, this time to the ECS of a homogeneous tissue (**Model 3**). In the ECS H_2_O_2_ can diffuse with diffusion coefficient *D_H_* or be consumed by a first-order process with rate constant *k*, which may account for permeation into the cells making up the tissue. For simplicity and to be conservative we neglect the permeation into the source cell. In spherical coordinates the dynamics of the H_2_O_2_ concentration at a radial distance *r* > *R* from the center of the cell is given by the following partial differential equation:

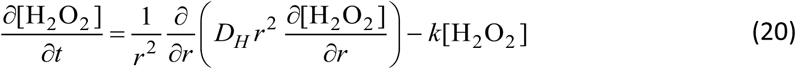

with boundary conditions

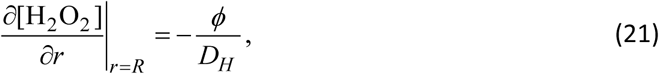

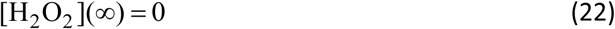

and steady-state solution

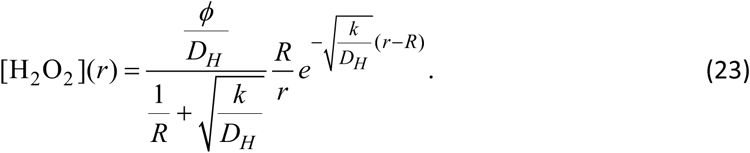

It follows from this solution that the maximal H_2_O_2_ concentration, which occurs at the cell surface (*r* = *R*), is:

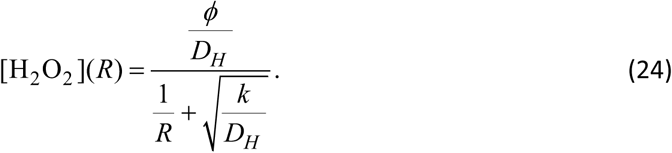

The concentration is highest in the *k* → 0 limit, where H_2_O_2_ simply diffuses away from the central cell and active clearance occurs at a negligible rate. Equation (23) then simplifies to

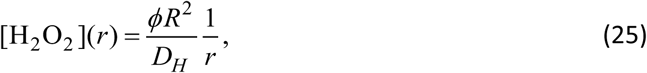

the H_2_O_2_ concentration at the cell surface becomes

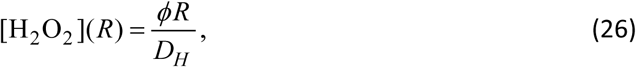

and decays to half this value at a distance *R* from the cell surface.

Setting again *R* = 4.2 μm, *ϕ* = 3.0 nmol dm^-2^s^-1^, *D_H_* = 1830 μm^2^s^-1^, equation (26) yields [H_2_O_2_](*R*) = 0.70 μM as the upper limit for the H_2_O_2_ concentration at the surface of a fully activated neutrophil embedded in a tissue of H_2_O_2_-impermeable cells and H_2_O_2_-unreactive ECS. That a model in which H₂O₂ diffuses with little impediment yields a far lower maximal concentration than the preceding ones makes two important points: diffusion is itself an efficient sink for locally generated H₂O₂, and local accumulation requires diffusion barriers such as the plasma membranes of neighboring cells.

To examine the effect of permeation into the tissue cells in this setting, we computed the apparent rate constant for clearance as follows. We idealized the embedding tissue as a reticulum of cubes with side *a* separated by a distance *d*. In this geometry, the fraction (*α*) of tissue volume occupied by the ECS and the ECS area/volume ratio (*Γ*) are thus

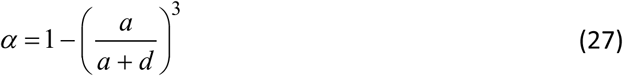

and

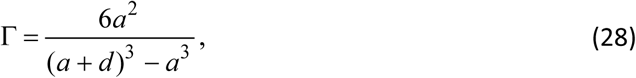

respectively. From this relationship one can readily relate the clearance rate constant of the tissue to the permeability constant of the cell membrane as

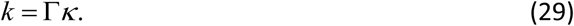

Considering *κ* = 10 µm s^−1^ and cubic cells with *a* = 15 µm side, from equations (28) and (29) one obtains *k* = 1.0x10^3^ s^−1^ and 19. s^−1^ for *d =* 20 nm and 1 µm separations between the cells, respectively.

In turn, Ledo *et al*. [12] determined *k* = 0.34 s^−1^ for the brain striatum of living rats, the low value being presumably due to the impermeability of myelinated cells. MPO released from activated neutrophils can substantially add to these clearance rates [29], and thus one must be mindful that these calculations are meant to provide upper bounds for the concentrations and decay length scales. They yield the radial H_2_O_2_ distributions and maximal concentrations shown in Figure 2A,B, respectively. The H_2_O_2_ concentrations decay to half their maximal values by 0.72, 2.4, 3.8 and 4.2 µm away of the cell membrane for *k* = 1.0x10^3^, 19., 0.34 and 0 s^−1^, respectively. Maximal concentrations are 0.17, 0.49, 0.66 and 0.7 µM, respectively.

**Figure 2.**
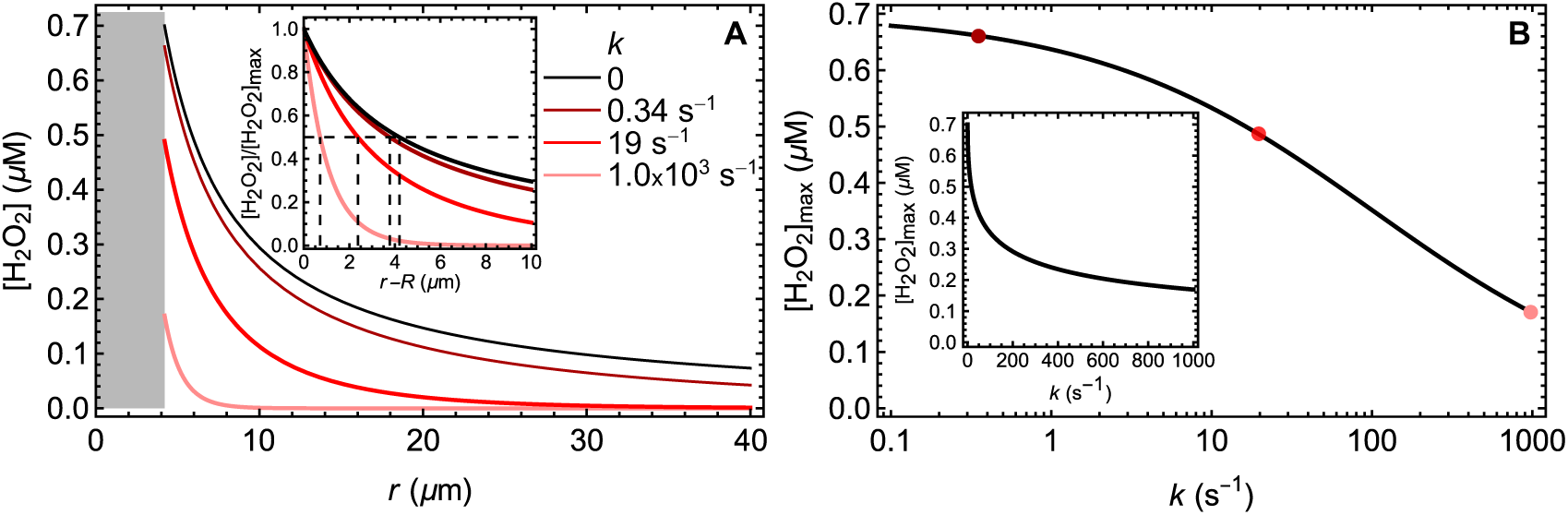
H_2_O_2_ distribution around a spherical activated neutrophil embedded in tissues of various clearance activities. (**A**) Concentration as function of the distance from the center of the cell. The inset shows the scaled concentrations as function of the distance from the cell surface, with the distances at which half-maximal concentrations obtain marked by vertical dashed lines. (**B**) Concentrations at the cell surface as function of the clearance rate constant. Note the logarithmic k scale. The inset shows the same function in Cartesian coordinates.

Treating the embedding tissue as a continuum as done in this model is admittedly a rough approximation. Nevertheless, these results highlight that (i) extracellular µM H_2_O_2_ concentrations are hardly attainable under physiological conditions when H_2_O_2_ is free to diffuse from the producing cells; (ii) the length scale of the gradients is not larger than the radius of the producing sites.

The situation in a tissue where H_2_O_2_ diffusion proceeds through the ECS interstices constrained by cells is intermediate between those in the models above, and so are the H_2_O_2_ concentrations. We highlight again that the area-specific production rate considered in the calculations above is an upper bound. However, the H_2_O_2_ concentrations in all the models above are directly proportional to the area-specific production rates, and therefore predicted concentrations can be straightforwardly scaled to more-physiological rates.

All the models above relied on the assumption that H_2_O_2_ is uniformly released throughout the plasma membrane of the source cell. However, as discussed in the Introduction, O_2_^•−^/H_2_O_2_ production is often localized at NOX/DUOX clusters in caveolae, lipid rafts or phagocytic cups. The dynamics and localization of O_2_^•−^ release from phagocytizing human neutrophils has been studied in sufficient detail to allow estimating area-specific O_2_^•−^/H_2_O_2_ release rates from phagocytic cups. In these cells, NOX2 is assembled already at the nascent cup of emerging phagosomes [115], and O_2_^•−^ production occurs within a minute of phagocytic stimulation [29]. O_2_^•−^ produced at the nascent cups is released to the ECS, but then the cups close and are internalized as phagosomes where the number of active NOX2 molecules continues to build up. Consequently, although total O_2_^•−^ production continues to increase over 10 – 20 min, release to the ECS peaks and wanes within 1 – 2 min of stimulation [116]. In the nascent cups, NOX2 forms clusters with an average 30 nm-radius, each containing about 10 NOX2 molecules [117]. NOX2 activity determinations based on a cell-free system, consisting of purified relipidated and reflavinated cytochrome b-559 and recombinant cytosolic components yielded a maximal rate of ∼320 O_2_^•−^ molecules/NOX2/s [118]. Accordingly, each 10-NOX2 cluster should produce a maximum of 3.2x10^3^ O_2_^•−^ molecules/s. However, the 6.4×10^6^ molecules/cell/s production computed by considering that each neutrophil carries ∼2000 NOX2 clusters at its surface [117] falls about one order of magnitude short of O_2_^•−^ release rates by activated neutrophils [29]. The most likely explanation for this discrepancy is that *in vivo* NOX2 can attain a substantially higher specific activity than determined in ref. [118]. We therefore adopted as reference a 10-fold higher rate, 3.2x10^4^ O_2_^•−^ molecules s^−1^ per average (30 nm radius) NOX2 cluster. Because all the concentrations reported below scale linearly with this rate, a reader who prefers the cell-free estimate [118] should divide them by 10. To estimate the concentrations and distribution of H_2_O_2_ from the instantaneous dismutation of the O_2_^•−^ released by a circular NOX2 cluster to the ECS between a neutrophil and another cell we considered the cylindrical geometry illustrated in Figure 1A (left). We assumed that both membranes have a *κ* = 10 µm s^−1^ permeability constant and that there are no barriers to H_2_O_2_ diffusion perpendicular to the plane of the membranes. The mathematical formulation of this model (**Model 4**) and the numerical methods applied for its integration are described in the Methods section.

Figure 3A shows the H_2_O_2_ distribution in the ECS for 30 nm and for 960 nm separation between the cells and a 30 nm NOX2 cluster radius. For a 30 nm separation between the cells, in the range of cell-cell adhesions, and an average *R*= 30 nm NOX2 cluster radius, the maximal H_2_O_2_ concentration is reached near the cluster’s center just below the source cell. Noably, it does not exceed 0.54 µM (Figure 3B). At the surface of the receptor cell the maximal concentration decreases to 0.47 µM. As the distance between both cells increases, the maximal concentration at the surface of the source cell converges asymptotically to a constant value ≍0.18 µM (Figure 3B,C). Because under physiological conditions *κ* << *D_H_* / *R* holds even for µm-scale *R*, this asymptotic concentration is mainly determined by a balance between production and diffusion and given by

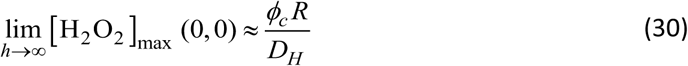

**Figure 3.**
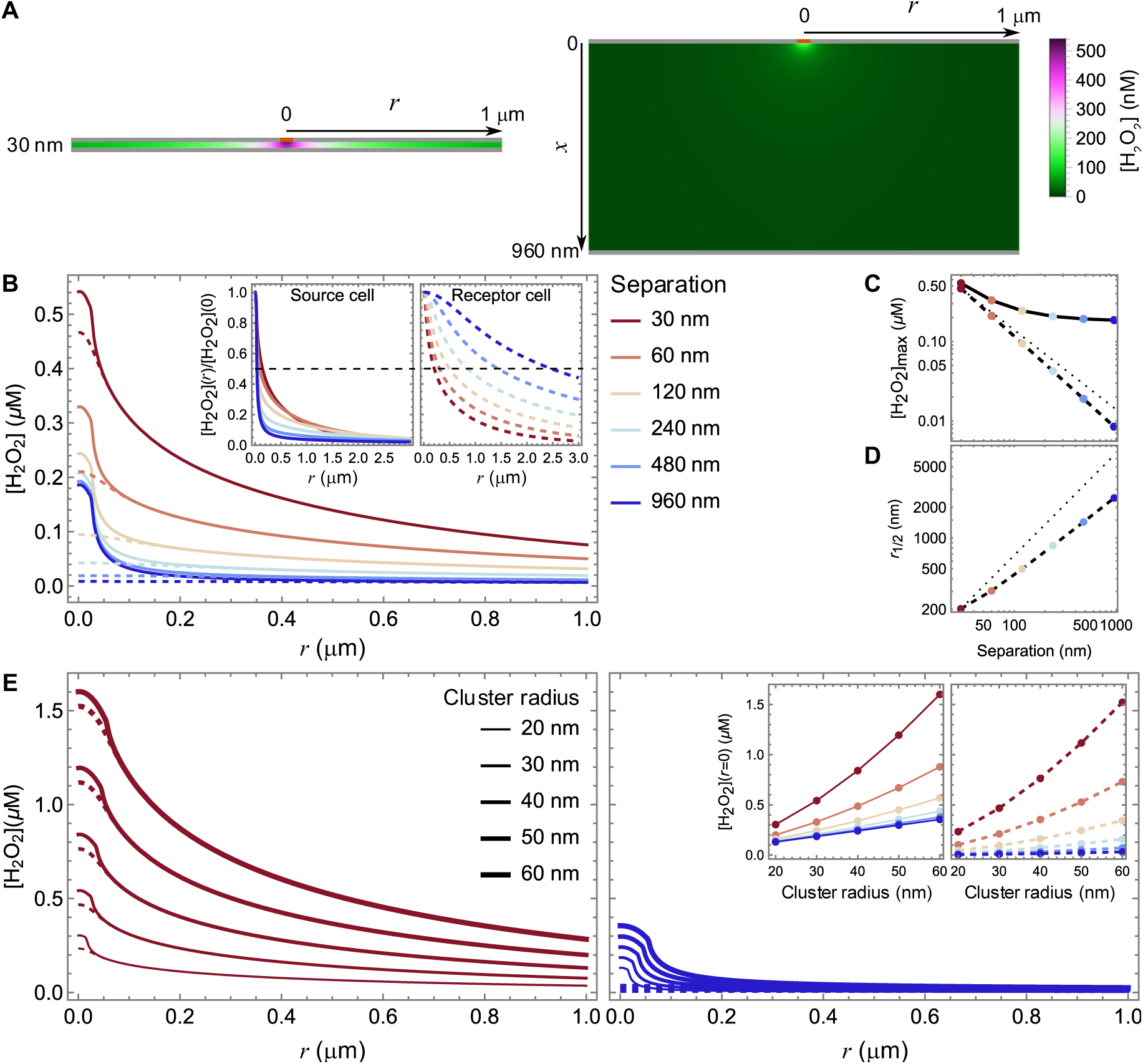
Distribution of H_2_O_2_ released from a circular NOX2 cluster to the ECS. (**A**) H_2_O_2_ distribution in the ECS for 30 nm (left) or 960 nm (right) separation between the source (top) and receptor (bottom) cells for a 30 nm radius cluster. The position of the cluster is marked in orange. (**B**) Radial H_2_O_2_ distribution near the surface of the source (solid lines) and receptor (dashed lines) cells for a 30 nm radius cluster and various separations between cells. The insets show the concentrations normalized by their values at or directly below the center of the cluster (r=0). The horizontal black dashed lines therein mark a 50% decrease from maximal values. (**C**) Maximum H_2_O_2_ concentrations near the surface of the source (solid lines) and receptor (dashed lines) cells for a 30 nm radius cluster, as function of the separation between the cells, highlighting the distinct dependence at both sites. Note the log scale. The lines connecting the points are guides for the eye. The thin dotted line marks a reciprocal relationship. (**D**) Radius at which the H_2_O_2_ concentration at the surface of the receptor cell declines to half-maximal values, as function of the separation between the cells. Note the log scale. The lines connecting the points are guides for the eye. The thin dotted line marks a proportional relationship. (**E**) Radial H_2_O_2_ distribution near the surface of the source (solid lines) and receptor (dashed lines) cells for various cluster radii and 30 nm (left) or 960 nm (right) separations between cells. The insets at the right show the maximal concentrations near the surface of the source (solid lines, left) and receptor (dashed lines, right) cells as function of the cluster radius for various separations between the cells. The lines in these insets are guides for the eye. A ϕ_c_= 5.7×10^6^ H_2_O_2_-equivalent molecules s^−1^µm^−1^ area-specific release rate by the clusters was assumed in all cases.

(see SI2). In turn, the maximal concentration at the surface of the receptor cell decreases almost reciprocally with the distance between the cells (slope of log-log plot of concentration at receptor cell as a function of distance is -1.2, see inset of Figure 3C). (We will analyze the reason for this scaling below.) Nevertheless, the *mean* concentration over the surface of the receptor cell decreases less steeply with the distance between the cells, and the same thus applies to the overall influx rate (Figure S1).

The maximal H_2_O_2_ concentration at the surface of both cells increases with the radius of the cluster (Figure 3E), as expected from the total release rate increasing proportionally to the cluster area. For short distances between the cells, the increase is supra-linear with the cluster radius (Figure 3E insets), though sublinear with the area (Figure S2). For larger inter-cellular distances, the increase with the radius becomes shallower and closer to linear.

In the radial direction, the H_2_O_2_ concentration near the membrane of the source cell decreases very steeply with distance from the cluster, and this decrease is steeper the greater the distance between the cells (Figure 3B, left hand inset). Thus, for a 30-nm radius cluster the concentration decreases to half-maximal values 145 nm and 36 nm away from the center of the cluster for 30 nm and 960 nm cell separations, respectively. The greater restriction to diffusion from the cluster and lower dilution when the ECS is narrow explain this trend. In contrast, near the membrane of the receptor cell the radial decrease of the concentration is less steep and becomes shallower as the distance between the cells increases (Figure 3B, right hand inset and Figure 3D). Thus, for a 30-nm radius cluster the concentration decreases to half-maximal values 210 nm and 2470 nm away from the center of the cluster for 30 nm and 960 nm cell separations, respectively.

The following approximate analysis helps understand both this trend and the approximate inverse proportionality of the maximal concentration at the receptor membrane on the cell-cell separation (*h*). For *h* >> *R* the cluster can be treated as a point source (*S*) releasing H_2_O_2_ at a rate *θ* to the ECS; and for physiological membrane permeabilities absorption by the source cell is negligible. Thus, absent any clearance or production in the ECS bulk, at steady state the total outward flux across any hemisphere of radius *ρ* centered at *S* (surface area 2 π *ρ*^2^) must equal *θ*. By Fick’s law this implies 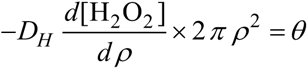, from which it follows that the concentration is inversely proportional to *ρ*, the distance from *S*. Within the surface of a parallel receptor membrane at distance *h* from the source, the distance of any point *P* to *S* is 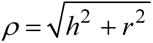, where *r* is the lateral distance from *P* to the point in the receptor membrane directly opposite *S*. Because the concentration is inversely proportional to *ρ*, it is maximal at the latter point and half-maximal at the lateral distance *r*_1/2_ that satisfies 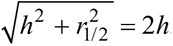. That is, 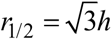. Because the receptor membrane constitutes a diffusion barrier in the axial direction, H_2_O_2_ spreads further in the radial direction than predicted form a spherical geometry, increasing *r*_1/2_. The linear relationship should still hold for *h*>>*R*, though. As *h* approaches *R*, the cluster size becomes the dominant effect.

Speculatively, even higher area-specific O_2_^•−^/H_2_O_2_ release rates may obtain near caveolae openings to the ECS. Large caveolae have a 50 nm radius [119] and 10 nm radius necks opening to the ECS [120], yielding an ≍ 4π (50 nm)^2^/(2π (10 nm)^2^)= 50-fold internal-to-neck surface area ratio. Hence, should all the caveola surface be covered with NOX2, the area-specific release rate (*ϕ*_Cav,max_= 2.9×10^8^ H_2_O_2_ equivalents s^−1^ µm^−2^) would be 50-fold higher than for a NOX2 cluster directly exposed to the ECS. Figure S3 shows the H_2_O_2_ distribution in the ECS near such a caveola neck. For adhering cells with a 30 nm separation, under this hypothetical extreme situation, at the surface of the source and receptor cells the H_2_O_2_ concentration could reach 6.5 µM and 3.7 µM, respectively (Figure S3).

In turn, once NOX-containing caveola or clathrin-coated pits [121] are internalized as closed endocytic vesicles (“redoxosomes” [36,122]), quite high lumenal H_2_O_2_ concentrations may be attained. A single maximally activated NOX2 complex releasing 1600 H_2_O_2_ equivalents/s inside an average 50 nm radius endocytic vesicle with a conservative *κ* = 10 μm s^−1^ membrane permeability yields a 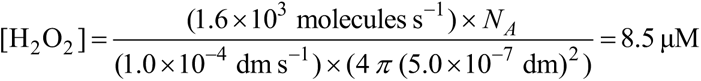 steady state concentration. This already substantially exceeds the maximal concentration in the ECS near a 30 nm-radius NOX2 cluster for 30 nm cell-cell separation. If such a cluster containing ≍10 NOX2 complexes is internalized, the concentration in the endosomal lumen might reach 85. µM. However, from equation (24) with *k* = *k_Se_*[Prdx]_T_ and values from Table 1 it follows that the H_2_O_2_ concentration at the cytosolic surface of the latter vesicles would be just 90. nM. Therefore, although the concentrations attained in the lumen can allow timely direct oxidation of moderately reactive protein thiols, this is not the case at the redoxosomes’ outer surface.

The H_2_O_2_ distribution and maximal concentrations in the ECS depend on the diffusion coefficient. Determinations of H_2_O_2_ diffusivity in the brain striatum of living rats [12] support a value in the range of that in bulk water (*D_H_*= 1.83×10^3^ µm^2^s^−1^ [96]), as adopted for all simulations above. However, the diffusivity may be substantially lower in some tissue environments. For instance, in a hydrogel with a viscosity matching that of the cytosol the diffusion coefficient is nearly 5-fold lower (*D_H_* = 3.7×10^2^ µm^2^s^−1^ [96]). Simulations as those shown in Figure 3 for diffusion coefficients in this range of values addressed their effect. The effect on the spatial distribution is relatively modest (Figure S4A), and the maximal concentrations at the surface of the source and receptor cells scale approximately as *D_H_*^−1/2^ (Figure S4B).

In neutrophils, NOX2 clusters are close enough that each point between the two cells receives H_2_O_2_ from multiple nearby clusters. To explore the consequences of this fact we drew on Figure 4A, left hand panel, of ref. [117] and on the results from the previous analysis to compute the “landscape” of H_2_O_2_ concentrations at the surface of an active neutrophil and of a nearby “bystander” cell. The original image represents fluorescently tagged NOX2 at the surface of a neutrophil-like cell (PLB-985 line) coated on poly-lysine. Analysis identifies 811 clusters distributed as shown in Figure 4A (black dots). Considering, as above, that each cluster has a 30 nm radius and yields a maximum of 16000 H_2_O_2_ molecules/s, one obtains a *ϕ* = 1.3 nmol dm^-2^s^-1^ mean area-specific release rate over the image surface delimited by the cell contour. While this is 2.4-fold less than the value considered in the first model, it is well within the range of experimentally determined release rates for activated neutrophils. For simplicity, we assume that the membrane of both the source and receptor cells are plane and that all the membranes of the cells laterally surrounding both these ones are, as well, permeable to H_2_O_2_ with permeability constant *κ* = 10 μm s^-1^. Applying the superposition principle to compute the distribution of H_2_O_2_ from the results of the previous single-cluster simulations yields the results shown in Figure 4A (see SI3 for image analysis and computation details).

**Figure 4.**
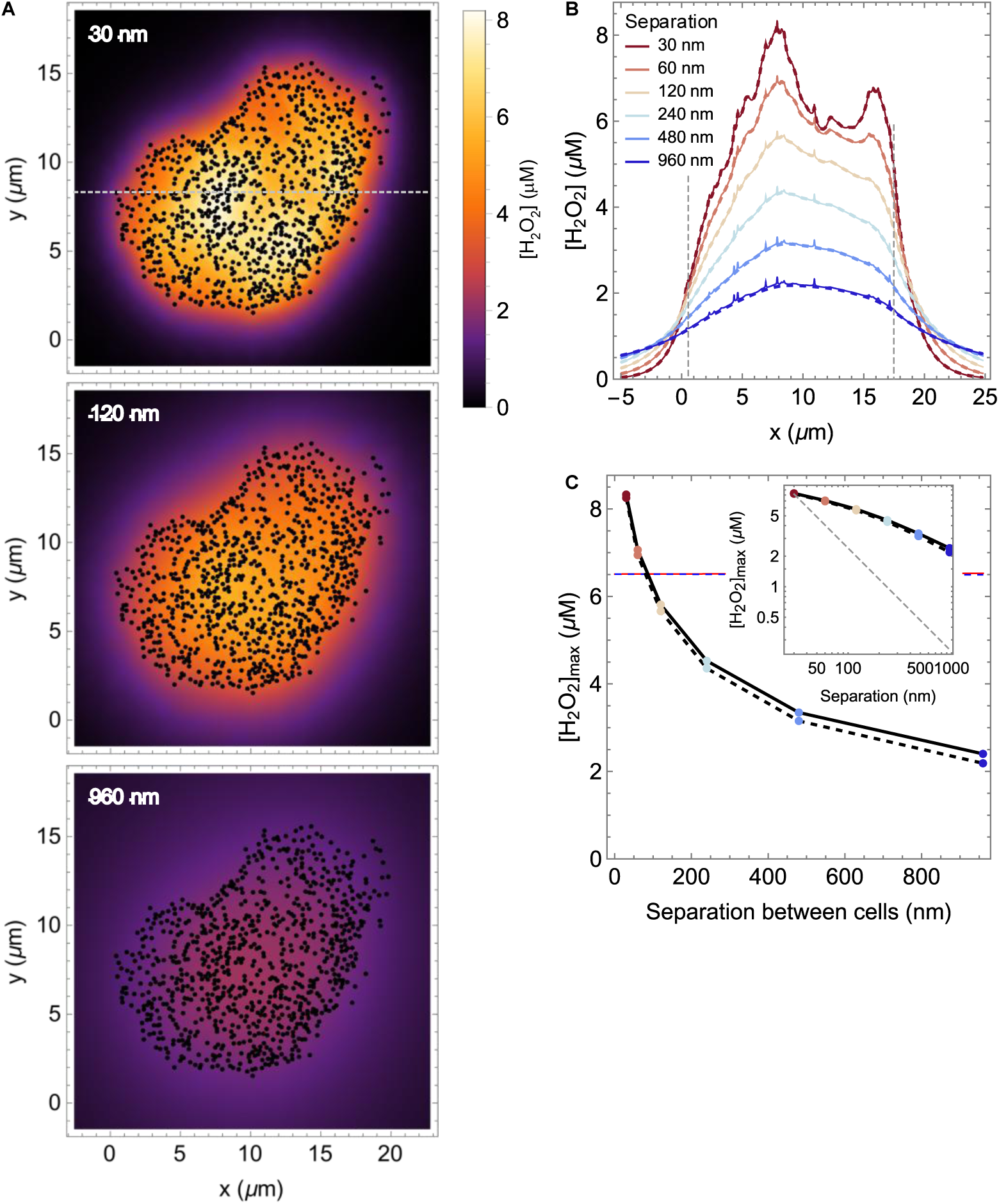
Computed distribution of H_2_O_2_ in the surface of an activated neutrophil and a bystander cell for various distances between them. (**A**) H_2_O_2_ concentrations at the surface of the receptor cells 30 nm, 120 nm and 960 nm apart from the neutrophil, considering 30-nm radius NOX2 clusters releasing H_2_O_2_ at maximal rates. The black spots mark the position of the NOX2 clusters, assumed to have 30 nm radius. The leftmost cluster lies at x=0. (**B**) H_2_O_2_ concentration along the dashed line in (A), which crosses the maximum, at the surface of the source (solid line) and the receptor (dashed) cells, for various separations between the cells. Small concentration spikes near the source cell reflect the proximity of NOX2 clusters. The vertical dashed lines mark the leftmost and rightmost NOX2 clusters along the line. (**C**) Maximal H_2_O_2_ concentration at the surface of the source (solid red line) and receptor (blue dashed line) cells. The inset shows the same in log-log coordinates, showing that the maximal concentration decreases less that reciprocally (thin gray dashed line) with the separation between cells over the considered range. The thin solid and dashed red lines in the main plot mark the H_2_O_2_ concentration at the surface of the source and receptor cells (respectively) according to equation (13) with ϕ set to the mean area-specific release rate computed from the cluster distribution in panel A as described in the main text.

Remarkably, the H_2_O_2_ concentrations attained near the surface of the source cell in this case are over one order of magnitude higher than those attained near an isolated 30-nm-radius NOX2 cluster. Moreover, the concentrations near the receptor cell are very similar to those near the source cell even for a 960 nm separation between these cells (Figure 4B), and they decrease less steeply with separation than the near-reciprocal dependence in the case of the isolated cluster (compare Figure 4C to Figure 3C). However, although the present situation approximates that of Model 1, they decrease much more steeply than predicted by Equation (13) (thin red and dashed blue lines in Figure 4C). Lateral leakage of the H_2_O_2_ to the ECS between non-H_2_O_2_-releasing cells near the neutrophil explains this steeper decrease: the higher the separation between source and receptor cells the higher the lifetime of the released H_2_O_2_ and the longer it can laterally diffuse away from the producing region. The steeper decline in H_2_O_2_ concentrations away from the neutrophil periphery at lower separations between source and receptor cell in Figure 4B (outside the vertical dashed lines) illustrates this point. Also important, even at the largest separation examined, the concentration halves within a few µm of the source-cell periphery, so little H₂O₂ reaches beyond the immediately laterally adjacent cells.

As a consequence of the broadening of the H_2_O_2_ distribution at the surface of the source cell with increasing cell-cell-separation (Figure 3B, right-hand inset), whereas for the small cell separations expected from adhering cells the concentration landscape between the two cells is quite rough, at higher separations it substantially smoothens (Figure 4B). The release of MPO to the ECS by the neutrophils may substantially decrease the concentrations above and somewhat roughen the landscape, but the simulations above capture the essential trends nevertheless.

All the models above rely on the assumption that H_2_O_2_ clearance by the cells is limited by the permeation barrier imposed by the plasma membrane. As argued in the Introduction, this is expected to be a good approximation under most physiological conditions. However, in the upper limits of H_2_O_2_ release, fast H_2_O_2_ inflow into the cytosol might deplete the antioxidant defenses near the inflow sites [15,33]. To scrutinize the extent and consequences of this phenomenon we extended Model 4 to consider reversible H_2_O_2_ permeation across the plasma membrane and H_2_O_2_ metabolism in the cytosol of both source and acceptor cells (Figure 1A) as described in the Numerical Models Section (**Model 5**). In these computationally demanding simulations, we consider just a 30 nm separation between the cells and a large, 60 nm radius, NOX2 cluster, but we examine a range of membrane permeabilities.

Even for this large NOX2 cluster and a permeability *κ* = 20 μm s^-1^, slightly higher than that of human erythrocyte membranes [60], the maximal area-specific H_2_O_2_ release rate considered above causes little oxidation of the intracellular redox pools (Figure 5C, F, G). The maximal concentrations of H_2_O_2_, Prdx-SOH, Prdx-SS, Trx-SS and GSSG in the cytosol of the receptor cell under this condition are just 10. nM, 2.2 µM, 0.15. µM, 86. nM and 0.97 nM, respectively (Figure 5B, D, F-L). Corresponding maxima in the source cell are at most a few percent higher, because H₂O₂ declines only slightly across the 30 nm of ECS separating the two cells. The gradients are slightly steeper in the axial direction than in the radial direction near the membrane (Figure 5H-L). This is because the gradients in the former direction track the more extended radial extracellular H_2_O_2_ gradient (Figure 5A), whereas in the latter direction they reflect mainly the high reductive activity in the cytosol. These gradients are determined by the balance between the half-lives and the diffusion constants of each species, as analyzed in further detail in ref. [33].

**Figure 5.**
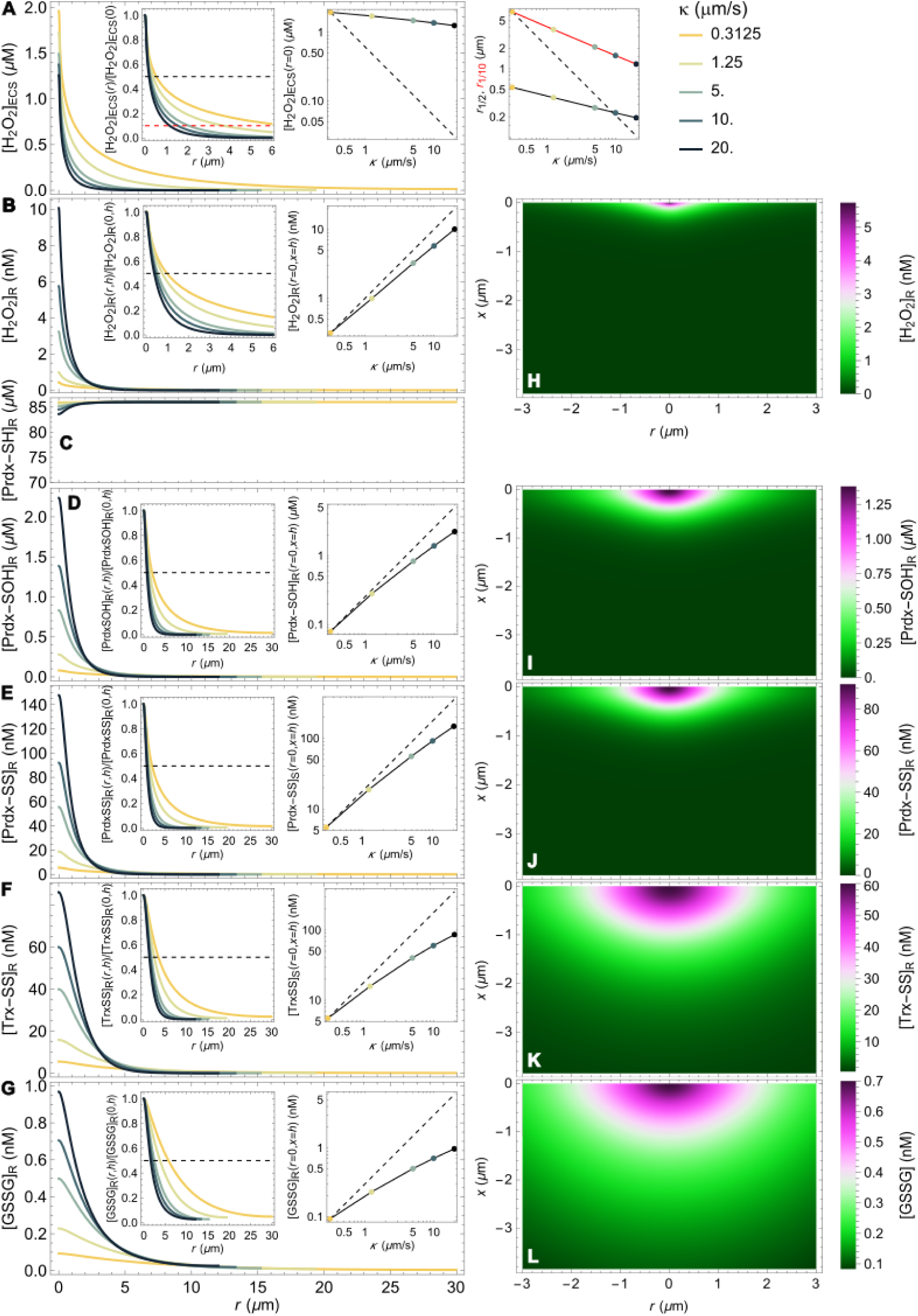
Distribution of H_2_O_2_ in the ECS and of the cytosolic redox pools in a receptor cell near a large (60 nm radius) maximally active NOX2 cluster at a 30 nm distant source cell. (**A-G**) Radial concentration distributions for a range of permeability constants of the membranes just above (**A**) or below (**B-G**) the receptor cell’s membrane cell. Note the expanded scale in C. Left-hand insets: normalized concentrations. Black and (in A) red dashed lines mark the half-maximal and 10%-maximal concentrations, respectively. Right-hand insets: dependence of the maximal concentrations on the permeability constant, in log-log coordinates. Dashed lines mark direct (inverse, in A) proportionality to the permeability constant, solid lines are guides for the eye. Right-hand panel in (**A**): dependence of the radial distance at which the H_2_O_2_ concentration declines to 50% (black line) or 10% (red line) of maximal on the permeability constant. Solid lines are guides for the eye. (**H-L**) Spatial distribution of the cytosolic redox pools in a segment of the receptor cell for κ= 10 μm/s. The light bands mark the regions where the concentrations are half-maximal.

The results above fully support the assumption of permeation-limited H_2_O_2_ clearance from the ECS in the previous models. The back-permeation of H_2_O_2_ from the source and receptor cells slightly increases in the maximal H_2_O_2_ in the ECS (compare Figure 5A curve for 60 nm radius cluster to Figure 3D curve for 10 µm/s permeability constant). Nevertheless, the maximal H_2_O_2_ concentration in the ECS decreases modestly with the increasing permeability constant, scaling approximately as [H_2_O_2_](0, 0) ∝ *κ* ^−0.11^ (Figure 5A, right hand inset). This is a shallower scaling than with the diffusion constant (Figure S4B), highlighting area-specific H_2_O_2_ release rate by the NOX2 cluster and diffusion as the main determinants of *maximal* H_2_O_2_ concentrations in this geometric setting. In turn, the effect of membrane permeability on the radial H_2_O_2_ distribution is somewhat stronger than on the maximal concentration, especially farther from the emitting cluster: the radii at which the H_2_O_2_ concentration declines to half and 1/10^th^ of the maximal value scale approximately as *r*_1/2_ ∝ *κ* ^−0.25^ and *r*_1/10_ ∝ *κ* ^−0.42^, respectively (Figure 5A, right hand panel).

Increased membrane permeability has two opposing effects on the intracellular concentrations of H_2_O_2_ and of the oxidized Prdx, Trx and glutathione forms: it decreases the H_2_O_2_ concentration in the ECS and it increases the inflow rate. For a fixed extracellular H_2_O_2_ concentration the latter increase is proportional to the permeability constant, and it is therefore the dominant effect. Thus, under this geometry, the maximal concentrations of H_2_O_2_, Prdx-SOH, Prdx-SS, Trx-SS, and GSSG in the receptor cell scale with the permeability constant with exponents 0.84, 0.80, 0.79, 0.66 and 0.56, respectively (Figure 5B, D-G, right hand insets). This holds under conditions where the cytosol’s capacity to reduce the oxidized pools everywhere substantially exceeds the H_2_O_2_ inflow rate.

Altogether, these results indicate that the O_2_^•−^/H_2_O_2_ production by a single large fully active NOX2 cluster is insufficient to substantially oxidize the Prdx pool of an adhering cell, even locally. However, as seen above, neutrophils display thousands of such clusters at their surface, the collective output of which may strongly elevate H_2_O_2_ concentrations in the ECS. To investigate how the intracellular redox pools behave under these conditions we used Model 6 where H_2_O_2_ is uniformly released throughout all the surface of the source cell that is in contact with the considered ECS segment (Figure 1B). Although the dimensions and release rates of the source cell are typical of a neutrophil, the simulations below are not meant to represent the dynamics of the redox pools in a neutrophil. This because they do not account, *e.g.*, for intracellular H_2_O_2_ release from internalized phagosomes or for the blocking of Prdx1/2-SS reduction by Trx1-SH [123] by a still unclear mechanism. Instead, they allow us to examine at once the effect of volume in cells with more typical cytosolic H_2_O_2_ release and cytosolic antioxidant defenses, as the receptor cell has twice the volume of the source cell and similar antioxidant defenses.

The H_2_O_2_ concentrations at the ECS-facing surfaces of both cells are similar too, given the small 30 nm separation between them. We considered the dynamics of O_2_^•−^/H_2_O_2_ release by neutrophils over the first 5 min after stimulation as characterized in ref. [116] (Figure 6A): the release rate increases over the first minute as NOX2 molecules in the phagocytic cups at the cell surface are activated, and then decreases over the next 4 min, presumably as the phagocytic cups close and are internalized. We considered that the maximal peak (*i. e.*, at 1 min) area-specific H_2_O_2_ release rate (*ϕ*_peak,max_= 12. nmol dm^−2^ s^−1^) is that of a maximally activated neutrophil concentrated at the area (in orange in Figure 1B) facing the considered ECS segment, and we ran simulations for a range of *ϕ*_peak_ values down to *ϕ*_peak,max_ /64.

**Figure 6.**
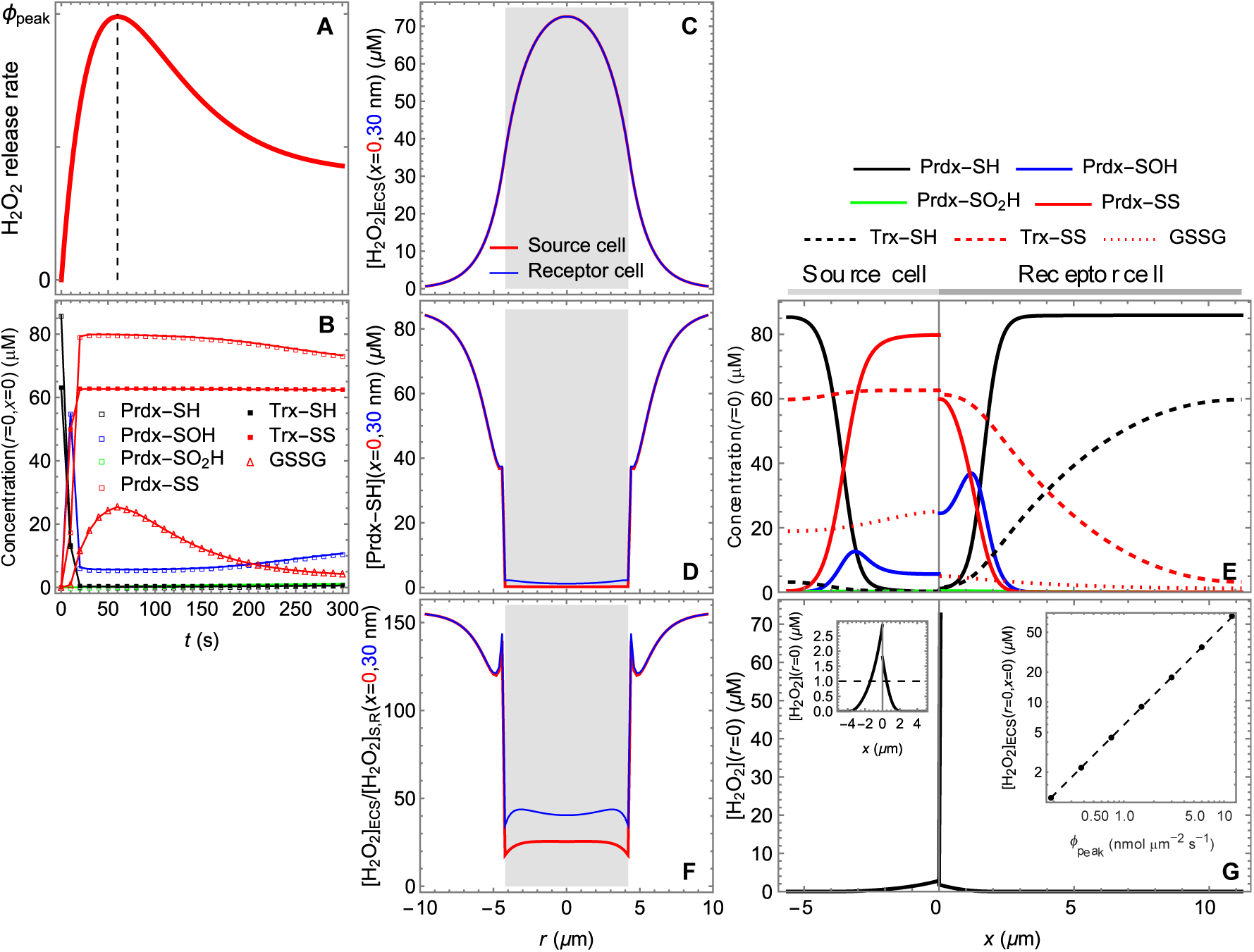
Distribution of H_2_O_2_ in the ECS and of the cytosolic redox pools in cells near a maximally active neutrophil. (Refer to Figure 1B for geometry.) (**A**) Time course of H_2_O_2_ release from synchronously activated neutrophils, assuming that all the O_2_^•−^ is instantaneously dismutated (replotted from [116]). The vertical dashed line marks the time t = 60 s at which release peaks. The remaining plots consider ϕ_peak_ = 12. nmol µm^−2^ s^−1^. (**B**) Time course of the concentrations of Prdx and Trx1 forms and of GSSG near the ECS-facing membrane of the source cell (**C**) Radial distribution of H_2_O_2_ in the ECS at the surface of the cells at the source and receptor sides a t = 60 s. The gray region marks range of r spanned by the source and receptor cells. (**D**) Radial distribution of Prdx-SH near the membranes of the cells at the source and receptor sides a t= 60 s. (**E**) Distribution of Prdx and Trx1 forms and of GSSG along the radial center of geometry at t= 60 s. (**F**) H_2_O_2_ concentration gradients across the membranes of source and receptor cells and their radial neighbors at t= 60 s. (**G**) H_2_O_2_ distribution along the axial direction at r= 0, t= 60 s. The right-hand inset shows the detail of the intracellular H_2_O_2_ distribution near the membranes. The left-hand inset shows the maximal H_2_O_2_ concentration in the ECS as function of the peak area-specific H_2_O_2_ release rate. Note the log-log scale. The dashed line marks a linear scaling.

The dynamics of the distribution of the various species is shown in Supplementary Movie 1 for *ϕ_peak_* = *ϕ*_peak,max_ and in Figure 6B. Except for Prdx-SO_2_H, which remains a very minor Prdx form in all the examined conditions, the dynamics of all the variables is fast relative to the variation in *ϕ*. Therefore, the oxidation of the Prdx, Trx and GSH pools peaks by ≍1 min (Figure 6B). So, below we focus on this time point.

At *ϕ*_peak,max_ the H_2_O_2_ concentration in the ECS reaches 70 µM, but falls to sub-µM values within 6 µm of the source cell, radially (Figure 6C). (The maximal concentration attained in these simulations considerably exceeds the ∼8.5 µM in Figure 4B for the same cell-cell separation because the mean area-specific H_2_O_2_ release rate is proportionally higher.) The former high concentrations overwhelm the capacity of the Trx1/TrxR system to reduce the 2-Cys Prdx near the ECS-facing membranes of both cells (Figure 6D). This effect is more pronounced in the smaller cell, given the lower total amount of Trx1 and overall cytosolic TrxR activity. However, the Prdx pools in both cells remain very reduced far from these membranes (Figure 6E), and the intracellular H_2_O_2_ concentrations decline from a few µM near the membrane to nM within 2 – 4 µm (Figure 6G, left-hand inset). Moreover, the GSH pool remains very reduced throughout, and the activity of the GPx/GSH/GSR system near the ECS-facing membranes is sufficient to ensure a 20-to 40-fold H_2_O_2_ gradient across these membranes (Figure 6F,G). Consequently, H_2_O_2_ clearance from the ECS remains permeation limited even in this condition, and the maximal H_2_O_2_ concentration still scales linearly with the area-specific H_2_O_2_ release rate (Figure 6G, right-hand inset). A different outcome with regards to the GSH pool would ensue should the modelled cells have as low a NADPH regeneration capacity as human erythrocytes, but in the latter cells the activity of catalase in the cytosol would suffice to keep a strong transmembrane H_2_O_2_ gradient [83].

Curiously, under these conditions the concentration of Prdx-SOH peaks at some distance from the ECS-facing membranes (Figure 6E, also observed in [33]). This is because Prdx-SOH formation requires both Prdx-SH and H_2_O_2_, and whereas the former reactant is depleted near these membranes the latter is depleted farther from them.

Again we highlight that these simulations address a very extreme scenario of very concentrated H_2_O_2_ release and, unrealistically for active neutrophils, absence of MPO activity in the ECS. Altogether, these results suggest that provided that the neutrophil output does not compromise the integrity of the plasma membranes the H_2_O_2_ they produce is unlikely to directly kill neighboring host cells.

## Discussion

Understanding the physiological extracellular concentrations and spatial distribution of H₂O₂ is essential for defining the constraints under which redox signaling operates in intact tissues. The present work used a set of reaction-diffusion models to delineate how geometry, membrane permeability, intracellular antioxidant capacity, and the localization of NOX/DUOX sources jointly determine extracellular H₂O₂ levels and distribution. Importantly, the concentrations attained in most of the results above set *upper bounds* for physiological situations. This is because these calculations rely on several assumptions that tend to inflate the H_2_O_2_ concentrations. Namely, that (i) O_2_^•−^ release rates are as for artificially stimulated neutrophils or clusters of maximally active NOX2, (ii) *all* this O_2_^•−^ instantaneously dismutates to 1/2 H_2_O_2_ and (iii) the latter is only cleared by diffusion over the ECS and permeation into cells. Losses through side reactions and cellular absorption of O_2_^•−^ and ECS H_2_O_2_ clearance by extracellular peroxidases are difficult to estimate. But these effects aside, the ECS H_2_O_2_ concentrations can be straightforwardly extrapolated to more-physiological release rates, as the former scale linearly with the latter under all the conditions here considered.

Our analyses show that µM-level H₂O₂ in the ECS is only attained under H_2_O_2_ release rates corresponding to maximally stimulated neutrophil output combined with confinement that minimizes diffusion away from production sites. The following results illustrate this point well and are also relevant for redox signaling by other cell types. NOX2 clusters as formed at the phagocytic cups of activated neutrophils likely yield the strongest localized O_2_^•−^ supply rates in human physiology when fully active. Yet, even with the above-mentioned favorable assumptions, H_2_O_2_ near a very large (60 nm radius) cluster hardly reaches 1.6 µM for intercellular distances typical of cell adhesion (Figure 3D). Highly localized concentrations up to ∼7 μM near the openings of hypothetically NOX-saturated caveolae to the ECS (Figure S3) remain a speculative possibility. And while the maximal concentration at the surface of an adhering receptor cell is in the same range, it declines approximately in the inverse proportion of cell-cell distance, reaching 4.3 nM at a 960 nm distance (Figure 3C). These maximal concentrations are modestly influenced by uncertainties about the permeability of the plasma membrane (Figure 5A) or H_2_O_2_‘s diffusion constant in the ECS (Figure S4).

These concentration ranges and their spatial distribution have implications for autocrine and paracrine signaling. Near the source cell, H_2_O_2_ is highly localized to a nanodomain extending a few 10’s to 120 nm radially beyond the NOX2 cluster, the more localized the greater the distance to the nearest cell. Cells in the neighborhood place diffusion barriers that enhance the local H_2_O_2_ concentration, but the maximum concentration decreases asymptotically with the cell-cell separation to a value in the 100’s nM (Figure 3C) that should still allow effective localized autocrine signaling [124]. In turn, at the surface of the receptor cell, H_2_O_2_ concentrations not only steeply decrease but also become more spatially distributed with increasing cell-cell separation. Thus, for adhering receptor cells H_2_O_2_ released from the NOX2 cluster is also confined to nanodomains. Juxtacrine signaling under these conditions is also highly localized intracellularly (Figure 5), given the small diffusion ranges of H_2_O_2_ and oxidized 2-Cys Prdx in the cytosol, as previously noted [15,33] (see also Figure 5H-J). This mode of signaling may be instrumental in the differential modulation of the cytoskeleton [125–127]. In turn, at µm cell-cell distances, as for cells embedded in extracellular matrix, the H_2_O_2_ domain at the surface of the receptor cell(s) spreads over a radius of several µm and may reach a few cells. It may thus allow near-cell paracrine signaling. Although low, the 10-nM-range concentrations at the receptor cell’s surface are spread over a much larger area, so the integrated influx is of the same order of magnitude as that driven by the higher but tightly localized concentrations of the preceding case (Figure S1). Results from an extension of the model in ref. [33] (Griffith, Araújo, Travasso, Salvador, manuscript in preparation) suggest that such concentrations can in principle drive Prdx2-mediated redox relays to transcription factors, if aided by scaffolds targeting both Prdx2 and target to the plasma membrane [e.g., 128].

Spatial heterogeneities in permeability in the cis- or transcellular neighborhood of NOX or DUOX clusters may influence the effectiveness of autocrine or juxtacrine signaling, respectively. The lipid composition of caveola / lipid rafts to which these clusters are usually associated renders them less permeable to H_2_O_2_ than the bulk of the plasma membrane [60]. This lower permeability detracts from autocrine signaling but enhances juxtacrine signaling by abating both the premature leakage of H_2_O_2_ from caveolae to the cytosol and the local competition of the source cell for released H_2_O_2_. In contrast, the presence of peroxiporins at the source cell’s membrane near NOX/DUOX clusters should favor autocrine over juxtacrine signaling. As a possible example, AQP3 has been shown to interact with NOX2 and to be required for H_2_O_2_-mediated NF-κB signaling in keratinocytes [61]. In turn, at the membrane of an adhering receptor cell, the presence of peroxiporins or clusters thereof within the H_2_O_2_ nanodomain’s range could strongly enhance juxtacrine signaling. Intracellularly, there is evidence of direct redox communication between the endoplasmic reticulum and mitochondria through contact sites between these organelles [129–131]. Whether extracellular redox synapses exist that promote the co-localization of NOX/DUOX clusters and peroxiporins across adhering cells is an interesting open question.

The large O_2_^•−^/H_2_O_2_ release rates of neutrophils are achieved through the activation of ∼2000 NOX2 clusters, which densely populate the plasma membrane. Even between adhering cells H_2_O_2_ can diffuse farther enough from the release sites that any point between the cells receives H_2_O_2_ from multiple clusters. Consequently, the local concentration is more reflective of the overall production over various clusters in their neighborhood than of that from the nearest cluster. This explains why substantially higher H_2_O_2_ concentrations – though still in the µM range – can be attained in this case (Figure 4). Though adhering receptor cells meet a rough H_2_O_2_ landscape at their surface, this landscape becomes quite smooth at intercellular distances beyond 100 nm (Figure 4B). However, these concentrations decline less steeply with cell separation than in the case of localized H_2_O_2_ release discussed in the previous paragraph (Figure 4C). In the radial direction, beyond the confines of the source cell the H_2_O_2_ concentration steeply declines within a few µm.

At such high H_2_O_2_ release rates, causing a substantial oxidative stress, the H_2_O_2_ influx from the ECS may strongly oxidize the Prdx and Trx pools near the release site. Of note, cell size can substantially influence the extent of Prdx-SH depletion and Prdx-SS accumulation (Figure 6E), as larger volume implies a greater amount of Prdx-SH available to react with inflowing H_2_O_2_ and of Trx-SH to reduce Prdx-SS. The local depletion of the Prdx-SH pool allows intracellular H_2_O_2_ to reach low-µM concentrations near the membrane facing this site (Figure 6D,E,G), declining to nM values within 2 – 4 µm. However, the transmembrane gradient remains quite substantial (Figure 6F,G). This happens because the GSH pool remains virtually intact, which allows GPx reaction to strongly outcompete H_2_O_2_ efflux.

Nevertheless, the inability of the released H_2_O_2_ to fully oxidize the cytosolic redox pools of adjacent cells with intact membranes does not imply that these cells can survive an oxidative burst. Neutrophils’ oxidative burst is associated with the ready release of myeloperoxidase (MPO) [29], which catalyzes the reaction of H_2_O_2_ with Cl^−^ to generate HOCl. While this reaction can substantially decrease the extracellular H_2_O_2_ concentrations [29], its HOCl product reacts rapidly with membrane proteins and lipids, forming chlorinated and cross-linked species that increase permeability and can lead to cell lysis [132]. Moreover, in inflammation neutrophils swarm, eventually forming dense clusters around sites of injury or infection, with the death of a limited number of neutrophils at such clusters acting as a swarming-enhancing positive feedback [133]. A recent preprint reports that the onset of swarming coincides with the onset of cell-cell coordinated H_2_O_2_ production by neutrophils [134]. However, in agreement with our results, H₂O₂ elevation in this circumstance was not associated with cell death, which occurred only hours after swarming onset [134]. The H_2_O_2_ concentrations that can be attained in the ECS of neutrophil clusters remains a topic for future analyses.

Experimental determinations of extracellular H_2_O_2_ concentrations and transport ranges remain scarce and uncertain. To our awareness, the following three are the quantitative studies that most directly address this issue. First, in the zebrafish tail wounding experimental model [1], epithelial cells at wound margins release DUOX-derived H_2_O_2_ at a rate sufficient to overwhelm the intracellular clearance capacity of cells within a ∼30 µm range and reach ∼5 µM extracellular H_2_O_2_ [9]. Moreover, in the neighborhood of the wound, H_2_O_2_ clearance from the ECS does not appear to be primarily permeation-limited [9], unlike the scenario here addressed in Model 6. That distinct outcome might be due to an exceptional H_2_O_2_ release rate by the epithelial cells. However, we are unaware of evidence that DUOXes have a specific activity comparable to that of NOXes or can accumulate at these cells’ surface as extensively as NOX2 in activated neutrophils. Thus, a limited intracellular H_2_O_2_ clearance capacity by the GPx/GSH/GSR system – besides the Prdx/Trx/TrxR system as previously noted [9] – may be an essential component of this phenomenon. The competition for NADPH by O_2_ reduction *via* DUOXes may play a role in attenuating the H_2_O_2_ clearance rate by both systems.

Second, using a plasma-membrane-anchored nanosensor, Hosogi *et al*. [135] report H_2_O_2_ concentrations approaching 10 µM at localized 700 nm-diameter spots on the surface of cultured A549 lung cancer cells. Maximal concentrations were lower in cell incubated with diphenyleneiodonium chloride, a flavoprotein inhibitor. In this system H_2_O_2_ is released to the aqueous medium overlying the cell layer. Therefore, these local H_2_O_2_ concentrations are most directly comparable to the asymptotic limit given by equation (30). Substituting the reference values of Table 1 into this equation and solving for *R* shows that a maximally activated NOX2 cluster would need an implausible 3.9 µm diameter for the H₂O₂ concentration at its center to reach 10 µM. This discrepancy might suggest that cells somehow can release H_2_O_2_ at much higher local area-specific rates than NOX2 clusters allow. However, the experimental protocol in ref. [135] required an incubation time of 60 min, over which some of the nanosensor particles may have been endocytosed. As discussed above, much higher H_2_O_2_ concentrations can be attained in endocytic vesicles’ lumen than in the extracellular space near NOX clusters.

Third, in the brain striatum of living rats H_2_O_2_ has a 2.2 s half-life, allowing it to diffuse over 100 µm in the extracellular space [12]. These half-life and diffusion distance are orders of magnitude longer than those predicted in this work for other tissues. As discussed elsewhere [13], the most likely explanation for the slow H_2_O_2_ clearance from the ECS in the brain is that myelination renders the cells virtually impermeable. Further experimental studies are needed to test this hypothesis, though.

## Concluding remarks

This work delineates how the interplay between NOX/DUOX source geometry, ECS microstructure, membrane permeability, and intracellular peroxidase systems constrains extracellular H₂O₂ concentrations and limits signaling ranges to nm –µm scales. The models establish that, even under upper-bound neutrophil release rates, physiologically realistic conditions support only sub- to low-µM H₂O₂ near source membranes and tens-of-nM levels at paracrine distances, highlighting localized autocrine and juxtacrine signaling as dominant modes of H₂O₂ action. ECS H₂O₂ concentrations much above ∼10 µM are difficult to reconcile with endogenous sources in intact tissues. These estimates provide quantitative guidance for designing experimental H₂O₂ manipulations and interpreting sensor readouts in intact tissues. The results also raise the question of whether specialized “extracellular redox synapses” transcellularly co-locating NOX/DUOX clusters with peroxiporins exist that optimize juxtacrine signaling. They also motivate systematic experimental probing of tissue-specific ECS architecture and membrane permeability to refine these bounds across organs and disease states.

## Acknowledgements

We thank Dr. Christine Winterbourn (University of Otago, New Zealand), Dr. Tobias Dansen (University Medical Center Utrecht, The Netherlands) and Wytze den Toom (University Medical Center Utrecht, The Netherlands) for helpful discussions about aspects related to O_2_^•-^/ H_2_O_2_ production by neutrophils and careful review of an early version of the manuscript.

## Declaration of generative AI and AI-assisted technologies in the manuscript preparation process

During the preparation of this work the authors used Anthropic’s Claude Opus 5 in order to improve writing style and check accuracy of references. After using this tool/service, the authors reviewed and edited the content as needed and take full responsibility for the content of the published article.

## Abbreviations

AQP: aquaporin
Cat: catalase
DUOX: dual oxidases
ECS: extracellular space
GPx: glutathione peroxidase
GSH: glutathione
GSR: glutathione reductase
GSSG: oxidized glutathione
MPO: myeloperoxidase
NOX: NADPH oxidase
PMA: phorbol 12-myristate 13-acetate
Prdx: peroxiredoxin
SOD3: superoxide dismutase 3
Trx: thioredoxin
TrxR: thioredoxin reductase.

## Footnotes

1 The term “H_2_O_2_ equivalent” here denotes the amount of H_2_O_2_ obtained if all the O_2_^•−^ released is dismutated.

