## Supplementary information for "Concentration limits and localization of hydrogen peroxide in the extracellular space of solid tissues"

### Supplementary Information for
### Concentration limits and localization of hydrogen peroxide in the extracellular space of solid tissues

Arthur Itacarambi[1], Marcos Gouveia[2], Rui D. M. Travasso[1*], Armindo Salvador[3,4,5**]

[1] CFisUC, Department of Physics, University of Coimbra, Rua Larga, 3004-516 Coimbra, Portugal
[2] Group of Numerical Methods in Engineering, Department of Mathematics, School of Civil Engineering, and CITEEC, University of A Coruña, Campus de Elviña s/n, 15008 A Coruña, Spain
[3] CNC – Centre for Neuroscience Cell Biology, University of Coimbra, Rua Larga Edificio FMUC, Piso 1, 3004–504 Coimbra, Portugal
[4] CiBB - Centre for Innovative Biomedicine and Biotechnology, University of Coimbra, Rua Larga Edificio FMUC, Piso 1, 3004–504 Coimbra, Portugal
[5] Coimbra Chemistry Center - Institute of Molecular Sciences (CQC-IMS), University of Coimbra, Rua Larga, 3004-535 Coimbra, Portugal

### 1. Equations for Models 5 and 6

Please refer to Figure 1 and Table 1 form the main text for the geometric setting and parameter meanings and reference values. The terms and equations in gray in the equations below are neglected in Model 5 because they contribute negligibly for the dynamics under physiological conditions. However, they are potentially important at the higher oxidative loads considered in Model 6.

$$\frac{\partial[\mathrm{H_2O_2}]}{\partial t} = D_H \left( \frac{\partial^2[\mathrm{H_2O_2}]}{\partial r^2} + \frac{1}{r}\frac{\partial[\mathrm{H_2O_2}]}{\partial r} + \frac{\partial^2[\mathrm{H_2O_2}]}{\partial x^2} \right)$$

$$\frac{\partial[\mathrm{H_2O_2}]_\mathrm{S}}{\partial t} = D_{H,cyt} \left( \frac{\partial^2[\mathrm{H_2O_2}]_\mathrm{S}}{\partial r^2} + \frac{1}{r}\frac{\partial[\mathrm{H_2O_2}]_\mathrm{S}}{\partial r} + \frac{\partial^2[\mathrm{H_2O_2}]_\mathrm{S}}{\partial x^2} \right) - k_\mathrm{Se}[\text{Prdx-S}^-]_\mathrm{S}[\mathrm{H_2O_2}]_\mathrm{S} -$$

$$- \frac{[\mathrm{GPx1}]\dfrac{[\mathrm{H_2O_2}]_\mathrm{S}}{\Phi_1}\dfrac{[\mathrm{GSH}]_\mathrm{S}}{\Phi_2}}{\dfrac{[\mathrm{H_2O_2}]_\mathrm{S}}{\Phi_1} + \dfrac{[\mathrm{GSH}]_\mathrm{S}}{\Phi_2}} - k_\mathrm{Si}[\text{Prdx-SO}^-]_\mathrm{S}[\mathrm{H_2O_2}]_\mathrm{S} - k_\mathrm{Cat}[\mathrm{H_2O_2}]_\mathrm{S}$$

$$\frac{\partial[\text{Prdx-S}^-]_\mathrm{S}}{\partial t} = D_\mathrm{P10} \left( \frac{\partial^2[\text{Prdx-S}^-]_\mathrm{S}}{\partial r^2} + \frac{1}{r}\frac{\partial[\text{Prdx-S}^-]_\mathrm{S}}{\partial r} + \frac{\partial^2[\text{Prdx-S}^-]_\mathrm{S}}{\partial x^2} \right) +$$

$$+ k_\mathrm{R}[\text{Prdx-SS}]_\mathrm{S}[\text{Trx-S}^-]_\mathrm{S} - k_\mathrm{Se}[\text{Prdx-S}^-]_\mathrm{S}[\mathrm{H_2O_2}]_\mathrm{S}$$

$$\frac{\partial[\text{Prdx-SO}^-]_\mathrm{S}}{\partial t} = D_\mathrm{P10} \left( \frac{\partial^2[\text{Prdx-SO}^-]_\mathrm{S}}{\partial r^2} + \frac{1}{r}\frac{\partial[\text{Prdx-SO}^-]_\mathrm{S}}{\partial r} + \frac{\partial^2[\text{Prdx-SO}^-]_\mathrm{S}}{\partial x^2} \right) +$$

$$+ k_\mathrm{Se}[\text{Prdx-S}^-]_\mathrm{S}[\mathrm{H_2O_2}]_\mathrm{S} - k_\mathrm{C}[\text{Prdx-SO}^-]_\mathrm{S} - k_\mathrm{Si}[\text{Prdx-SO}^-]_\mathrm{S}[\mathrm{H_2O_2}]_\mathrm{S}$$

$$\frac{\partial[\text{Prdx-SO}_2^-]_\mathrm{S}}{\partial t} = D_\mathrm{P10} \left( \frac{\partial^2[\text{Prdx-SO}_2^-]_\mathrm{S}}{\partial r^2} + \frac{1}{r}\frac{\partial[\text{Prdx-SO}_2^-]_\mathrm{S}}{\partial r} + \frac{\partial^2[\text{Prdx-SO}_2^-]_\mathrm{S}}{\partial x^2} \right) +$$

$$+ k_\mathrm{Si}[\text{Prdx-SO}^-]_\mathrm{S}[\mathrm{H_2O_2}]_\mathrm{S} - k_\mathrm{Srx}[\text{Prdx-SO}_2^-]_\mathrm{S}$$

$$\frac{\partial[\text{Prdx-SS}]_\mathrm{S}}{\partial t} = D_\mathrm{P10} \left( \frac{\partial^2[\text{Prdx-SS}]_\mathrm{S}}{\partial r^2} + \frac{1}{r}\frac{\partial[\text{Prdx-SS}]_\mathrm{S}}{\partial r} + \frac{\partial^2[\text{Prdx-SS}]_\mathrm{S}}{\partial x^2} \right) + k_\mathrm{C}[\text{Prdx-SO}^-]_\mathrm{S} -$$

$$- k_\mathrm{R}[\text{Prdx-SS}]_\mathrm{S}[\text{Trx-S}^-]_\mathrm{S}$$

$$\frac{\partial[\text{Trx-S}^-]_\mathrm{S}}{\partial t} = D_\mathrm{Trx} \left( \frac{\partial^2[\text{Trx-S}^-]_\mathrm{S}}{\partial r^2} + \frac{1}{r}\frac{\partial[\text{Trx-S}^-]_\mathrm{S}}{\partial r} + \frac{\partial^2[\text{Trx-S}^-]_\mathrm{S}}{\partial x^2} \right) + \frac{V_\mathrm{TrxR}[\text{Trx-SS}]_\mathrm{S}}{K_\mathrm{M,Trx\text{-}SS} + [\text{Trx-SS}]_\mathrm{S}} -$$

$$- k_\mathrm{R}[\text{Prdx-SS}]_\mathrm{S}[\text{Trx-S}^-]_\mathrm{S}$$

$$\frac{\partial[\text{Trx-SS}]_\mathrm{S}}{\partial t} = D_\mathrm{Trx} \left( \frac{\partial^2[\text{Trx-SS}]_\mathrm{S}}{\partial r^2} + \frac{1}{r}\frac{\partial[\text{Trx-SS}]_\mathrm{S}}{\partial r} + \frac{\partial^2[\text{Trx-SS}]_\mathrm{S}}{\partial x^2} \right) + k_\mathrm{R}[\text{Prdx-SS}]_\mathrm{S}[\text{Trx-S}^-]_\mathrm{S} -$$

$$- \frac{V_\mathrm{Max}[\text{Trx-SS}]_\mathrm{S}}{K_\mathrm{M,Trx\text{-}SS} + [\text{Trx-SS}]_\mathrm{S}}$$

$$\frac{\partial[\mathrm{GSH}]_\mathrm{S}}{\partial t}=D_\mathrm{Trx}\left(\frac{\partial^2[\mathrm{GSH}]_\mathrm{S}}{\partial r^2}+\frac{1}{r}\frac{\partial[\mathrm{GSH}]_\mathrm{S}}{\partial r}+\frac{\partial^2[\mathrm{GSH}]_\mathrm{S}}{\partial x^2}\right)+2\frac{V_\mathrm{GSR}\dfrac{[\mathrm{GSSG}]_\mathrm{S}}{K_\mathrm{M,GSSG}}}{1+\dfrac{[\mathrm{GSSG}]_\mathrm{S}}{K_\mathrm{M,GSSG}}}-$$

$$-2\frac{[\mathrm{GPx1}]\dfrac{[\mathrm{H_2O_2}]_\mathrm{S}}{\Phi_1}\dfrac{[\mathrm{GSH}]_\mathrm{S}}{\Phi_2}}{\dfrac{[\mathrm{H_2O_2}]_\mathrm{S}}{\Phi_1}+\dfrac{[\mathrm{GSH}]_\mathrm{S}}{\Phi_2}}$$

$$\frac{\partial[\mathrm{GSSG}]_\mathrm{S}}{\partial t}=D_\mathrm{Trx}\left(\frac{\partial^2[\mathrm{GSSG}]_\mathrm{S}}{\partial r^2}+\frac{1}{r}\frac{\partial[\mathrm{GSSG}]_\mathrm{S}}{\partial r}+\frac{\partial^2[\mathrm{GSSG}]_\mathrm{S}}{\partial x^2}\right)+\frac{[\mathrm{GPx1}]\dfrac{[\mathrm{H_2O_2}]_\mathrm{S}}{\Phi_1}\dfrac{[\mathrm{GSH}]_\mathrm{S}}{\Phi_2}}{\dfrac{[\mathrm{H_2O_2}]_\mathrm{S}}{\Phi_1}+\dfrac{[\mathrm{GSH}]_\mathrm{S}}{\Phi_2}}-$$

$$-\frac{V_\mathrm{GSR}\dfrac{[\mathrm{GSSG}]_\mathrm{S}}{K_\mathrm{M,GSSG}}}{1+\dfrac{[\mathrm{GSSG}]_\mathrm{S}}{K_\mathrm{M,GSSG}}}$$

$$\frac{\partial[\mathrm{H_2O_2}]_\mathrm{R}}{\partial t}=D_{H,cyt}\left(\frac{\partial^2[\mathrm{H_2O_2}]_\mathrm{R}}{\partial r^2}+\frac{1}{r}\frac{\partial[\mathrm{H_2O_2}]_\mathrm{R}}{\partial r}+\frac{\partial^2[\mathrm{H_2O_2}]_\mathrm{R}}{\partial x^2}\right)-k_\mathrm{Se}[\mathrm{Prdx\text{-}S^-}]_\mathrm{R}[\mathrm{H_2O_2}]_\mathrm{R}-$$

$$-\frac{[\mathrm{GPx1}]\dfrac{[\mathrm{H_2O_2}]_\mathrm{R}}{\Phi_1}\dfrac{[\mathrm{GSH}]_\mathrm{R}}{\Phi_2}}{\dfrac{[\mathrm{H_2O_2}]_\mathrm{R}}{\Phi_1}+\dfrac{[\mathrm{GSH}]_\mathrm{R}}{\Phi_2}}-k_\mathrm{Si}[\mathrm{Prdx\text{-}SO^-}]_\mathrm{R}[\mathrm{H_2O_2}]_\mathrm{R}-k_\mathrm{Cat}[\mathrm{H_2O_2}]_\mathrm{R}$$

$$\frac{\partial[\mathrm{Prdx\text{-}S^-}]_\mathrm{R}}{\partial t}=D_\mathrm{P10}\left(\frac{\partial^2[\mathrm{Prdx\text{-}S^-}]_\mathrm{R}}{\partial r^2}+\frac{1}{r}\frac{\partial[\mathrm{Prdx\text{-}S^-}]_\mathrm{R}}{\partial r}+\frac{\partial^2[\mathrm{Prdx\text{-}S^-}]_\mathrm{R}}{\partial x^2}\right)+$$

$$+k_\mathrm{R}[\mathrm{Prdx\text{-}SS}]_\mathrm{R}[\mathrm{Trx\text{-}S^-}]_\mathrm{R}-k_\mathrm{Se}[\mathrm{Prdx\text{-}S^-}]_\mathrm{R}[\mathrm{H_2O_2}]_\mathrm{R}$$

$$\frac{\partial[\mathrm{Prdx\text{-}SO^-}]_\mathrm{R}}{\partial t}=D_\mathrm{P10}\left(\frac{\partial^2[\mathrm{Prdx\text{-}SO^-}]_\mathrm{R}}{\partial r^2}+\frac{1}{r}\frac{\partial[\mathrm{Prdx\text{-}SO^-}]_\mathrm{R}}{\partial r}+\frac{\partial^2[\mathrm{Prdx\text{-}SO^-}]_\mathrm{R}}{\partial x^2}\right)+$$

$$+k_\mathrm{Se}[\mathrm{Prdx\text{-}S^-}]_\mathrm{R}[\mathrm{H_2O_2}]_\mathrm{R}-k_\mathrm{C}[\mathrm{Prdx\text{-}SO^-}]_\mathrm{R}-k_\mathrm{Si}[\mathrm{Prdx\text{-}SO^-}]_\mathrm{R}[\mathrm{H_2O_2}]_\mathrm{R}$$

$$\frac{\partial[\mathrm{Prdx\text{-}SO_2^-}]_\mathrm{R}}{\partial t}=D_\mathrm{P10}\left(\frac{\partial^2[\mathrm{Prdx\text{-}SO_2^-}]_\mathrm{R}}{\partial r^2}+\frac{1}{r}\frac{\partial[\mathrm{Prdx\text{-}SO_2^-}]_\mathrm{R}}{\partial r}+\frac{\partial^2[\mathrm{Prdx\text{-}SO_2^-}]_\mathrm{R}}{\partial x^2}\right)+$$

$$+k_\mathrm{Si}[\mathrm{Prdx\text{-}SO^-}]_\mathrm{R}[\mathrm{H_2O_2}]_\mathrm{R}-k_\mathrm{Srx}[\mathrm{Prdx\text{-}SO_2^-}]_\mathrm{R}$$

$$\frac{\partial[\mathrm{Prdx\text{-}SS}]_\mathrm{R}}{\partial t}=D_\mathrm{P10}\left(\frac{\partial^2[\mathrm{Prdx\text{-}SS}]_\mathrm{R}}{\partial r^2}+\frac{1}{r}\frac{\partial[\mathrm{Prdx\text{-}SS}]_\mathrm{R}}{\partial r}+\frac{\partial^2[\mathrm{Prdx\text{-}SS}]_\mathrm{R}}{\partial x^2}\right)+k_\mathrm{C}[\mathrm{Prdx\text{-}SO^-}]_\mathrm{R}-$$

$$-k_\mathrm{R}[\mathrm{Prdx\text{-}SS}]_\mathrm{R}[\mathrm{Trx\text{-}S^-}]_\mathrm{R}$$

$$\frac{\partial[\mathrm{Trx\text{-}S^-}]_\mathrm{R}}{\partial t}=D_\mathrm{Trx}\left(\frac{\partial^2[\mathrm{Trx\text{-}S^-}]_\mathrm{R}}{\partial r^2}+\frac{1}{r}\frac{\partial[\mathrm{Trx\text{-}S^-}]_\mathrm{R}}{\partial r}+\frac{\partial^2[\mathrm{Trx\text{-}S^-}]_\mathrm{R}}{\partial x^2}\right)+\frac{V_\mathrm{TrxR}[\mathrm{Trx\text{-}SS}]_\mathrm{R}}{K_\mathrm{M,Trx\text{-}SS}+[\mathrm{Trx\text{-}SS}]_\mathrm{R}}-$$

$$-k_\mathrm{R}[\mathrm{Prdx\text{-}SS}]_\mathrm{R}[\mathrm{Trx\text{-}S^-}]_\mathrm{R}$$

$$\frac{\partial[\text{Trx-SS}]_\text{R}}{\partial t}=D_\text{Trx}\left(\frac{\partial^2[\text{Trx-SS}]_\text{R}}{\partial r^2}+\frac{1}{r}\frac{\partial[\text{Trx-SS}]_\text{R}}{\partial r}+\frac{\partial^2[\text{Trx-SS}]_\text{R}}{\partial x^2}\right)+k_\text{R}[\text{Prdx-SS}]_\text{R}[\text{Trx-S}^-]_\text{R}-$$

$$-\frac{V_\text{Max}[\text{Trx-SS}]_\text{R}}{K_\text{M,Trx-SS}+[\text{Trx-SS}]_\text{R}}$$

$$\frac{\partial[\text{GSH}]_\text{R}}{\partial t}=D_\text{Trx}\left(\frac{\partial^2[\text{GSH}]_\text{R}}{\partial r^2}+\frac{1}{r}\frac{\partial[\text{GSH}]_\text{R}}{\partial r}+\frac{\partial^2[\text{GSH}]_\text{R}}{\partial x^2}\right)+2\frac{V_\text{GSR}\dfrac{[\text{GSSG}]_\text{R}}{K_\text{M,GSSG}}}{1+\dfrac{[\text{GSSG}]_\text{R}}{K_\text{M,GSSG}}}-$$

$$-2\frac{[\text{GPx1}]\dfrac{[\text{H}_2\text{O}_2]_\text{R}}{\Phi_1}\dfrac{[\text{GSH}]_\text{R}}{\Phi_2}}{\dfrac{[\text{H}_2\text{O}_2]_\text{R}}{\Phi_1}+\dfrac{[\text{GSH}]_\text{R}}{\Phi_2}}$$

$$\frac{\partial[\text{GSSG}]_\text{R}}{\partial t}=D_\text{Trx}\left(\frac{\partial^2[\text{GSSG}]_\text{R}}{\partial r^2}+\frac{1}{r}\frac{\partial[\text{GSSG}]_\text{R}}{\partial r}+\frac{\partial^2[\text{GSSG}]_\text{R}}{\partial x^2}\right)+\frac{[\text{GPx1}]\dfrac{[\text{H}_2\text{O}_2]_\text{R}}{\Phi_1}\dfrac{[\text{GSH}]_\text{R}}{\Phi_2}}{\dfrac{[\text{H}_2\text{O}_2]_\text{R}}{\Phi_1}+\dfrac{[\text{GSH}]_\text{R}}{\Phi_2}}-$$

$$-\frac{V_\text{GSR}\dfrac{[\text{GSSG}]_\text{R}}{K_\text{M,GSSG}}}{1+\dfrac{[\text{GSSG}]_\text{R}}{K_\text{M,GSSG}}}$$

The boundary conditions for the $H_2O_2$ concentrations in the ECS, in the <u>S</u>ource cell and in the <u>R</u>eceptor cell are:

$$\left.\frac{\partial[\text{H}_2\text{O}_2]}{\partial x}\right|_{x=0,r\le R}=\frac{\kappa([\text{H}_2\text{O}_2](x=0)-[\text{H}_2\text{O}_2]_\text{S}(x=0))-\phi_c}{D_H}, \quad (1)$$

$$\left.\frac{\partial[\text{H}_2\text{O}_2]}{\partial x}\right|_{x=0,r>R}=\frac{\kappa}{D_H}\big([\text{H}_2\text{O}_2](x=0)-[\text{H}_2\text{O}_2]_\text{S}(x=0)\big), \quad (2)$$

$$\left.\frac{\partial[\text{H}_2\text{O}_2]}{\partial x}\right|_{x=h}=-\frac{\kappa}{D_H}\big([\text{H}_2\text{O}_2](x=h)-[\text{H}_2\text{O}_2]_\text{R}(x=h)\big), \quad (3)$$

$$\left.\frac{\partial[\text{H}_2\text{O}_2]}{\partial r}\right|_{r=0}=0, \quad (4)$$

$$\left.\frac{\partial[\text{H}_2\text{O}_2]}{\partial r}\right|_{r\to\infty}=0, \quad (5)$$

$$\left.\frac{\partial[\text{H}_2\text{O}_2]_\text{S}}{\partial x}\right|_{x=0}=\frac{\kappa}{D_{H,cyt}}\big([\text{H}_2\text{O}_2](x=0)-[\text{H}_2\text{O}_2]_\text{S}(x=0)\big), \quad (6)$$

$$\left.\frac{\partial[\mathrm{H_2O_2}]_\mathrm{S}}{\partial x}\right|_{x=-h_\mathrm{S}}=0, \tag{7}$$

$$\left.\frac{\partial[\mathrm{H_2O_2}]_\mathrm{S}}{\partial r}\right|_{r=0}=0, \tag{8}$$

$$\left.\frac{\partial[\mathrm{H_2O_2}]_\mathrm{S}}{\partial r}\right|_{r\to\infty}=0, \tag{9}$$

$$\left.\frac{\partial[\mathrm{H_2O_2}]_\mathrm{R}}{\partial x}\right|_{x=h}=\frac{\kappa}{D_{H,cyt}}\left([\mathrm{H_2O_2}]_\mathrm{R}(x=h)-[\mathrm{H_2O_2}](x=h)\right), \tag{10}$$

$$\left.\frac{\partial[\mathrm{H_2O_2}]_\mathrm{R}}{\partial x}\right|_{x=h+h_\mathrm{R}}=0, \tag{11}$$

$$\left.\frac{\partial[\mathrm{H_2O_2}]_\mathrm{R}}{\partial r}\right|_{r=0}=0, \tag{12}$$

$$\left.\frac{\partial[\mathrm{H_2O_2}]_\mathrm{R}}{\partial r}\right|_{r\to\infty}=0. \tag{13}$$

These boundary conditions are common to Models 5 and 6, but for Model 6 we consider the following additional boundary conditions for $H_2O_2$:

$$\left.\frac{\partial[\mathrm{H_2O_2}]_\mathrm{S}}{\partial r}\right|_{r=R}=\left.\frac{\partial[\mathrm{H_2O_2}]_\mathrm{OS}}{\partial r}\right|_{r=R}=\frac{\kappa/2}{D_{H,cyt}}\left([\mathrm{H_2O_2}]_\mathrm{OS}(r=R)-[\mathrm{H_2O_2}]_\mathrm{S}(r=R)\right), \tag{14}$$

$$\left.\frac{\partial[\mathrm{H_2O_2}]_\mathrm{R}}{\partial r}\right|_{r=R}=\left.\frac{\partial[\mathrm{H_2O_2}]_\mathrm{OR}}{\partial r}\right|_{r=R}=\frac{\kappa/2}{D_{H,cyt}}\left([\mathrm{H_2O_2}]_\mathrm{OR}(r=R)-[\mathrm{H_2O_2}]_\mathrm{R}(r=R)\right), \tag{15}$$

where the “OS” and “OR” denote the outer “cells” surrounding the central source and receptor cell. In turn, we consider no-flux conditions at all boundaries for the other species’ concentrations in the cytosol of the source and receptor cells:

$$\left.\frac{\partial X_\mathrm{S}}{\partial x}\right|_{x=0}=\left.\frac{\partial X_\mathrm{S}}{\partial x}\right|_{x=-h_\mathrm{S}}=\left.\frac{\partial X_\mathrm{S}}{\partial r}\right|_{r=0}=\left.\frac{\partial X_\mathrm{(O)S}}{\partial r}\right|_{r\to\infty}=0, \tag{16}$$

$$\left.\frac{\partial X_\mathrm{R}}{\partial x}\right|_{x=h}=\left.\frac{\partial X_\mathrm{R}}{\partial x}\right|_{x=h+h_\mathrm{S}}=\left.\frac{\partial X_\mathrm{R}}{\partial r}\right|_{r=0}=\left.\frac{\partial X_\mathrm{(O)R}}{\partial r}\right|_{r\to\infty}=0, \tag{17}$$

for $X \in \{[\text{Prdx-S}^-], [\text{Prdx-SO}^-], [\text{Prdx-SO}_2^-], [\text{Prdx-SS}], [\text{Trx-S}^-], [\text{Trx-SS}], [\text{GSH}], [\text{GSSG}]\}$.

For Model 6 the following additional boundary conditions apply:

$$\left.\frac{\partial X_{\text{S}}}{\partial r}\right|_{r=R} = \left.\frac{\partial X_{\text{OS}}}{\partial r}\right|_{r=R} = 0, \tag{18}$$

$$\left.\frac{\partial X_{\text{R}}}{\partial r}\right|_{r=R} = \left.\frac{\partial X_{\text{OR}}}{\partial r}\right|_{r=R} = 0. \tag{19}$$

#### 2. Derivation of Equation 30 from the main text

Neglecting the absorption of $H_2O_2$ by permeation into the cell, we consider a circular cluster of radius $R$ on an infinite planar, otherwise impermeable, membrane that releases $H_2O_2$ at a constant area-specific rate $\phi$. $H_2O_2$ diffuses freely into the adjacent semi-infinite bulk medium with diffusion coefficient $D_H$. We seek the steady-state concentration $[\text{H}_2\text{O}_2](r, x)$, where $x \geq 0$ is the distance from the membrane and $r$ the radial distance from the cluster center, and in particular its maximal value, attained at the center of the patch $(r = 0, x = 0)$.

In the bulk, mass conservation with no reaction term reduces to Laplace's equation in cylindrical coordinates,

$$\frac{\partial[\text{H}_2\text{O}_2]}{\partial t} = D_H\left(\frac{\partial^2[\text{H}_2\text{O}_2]}{\partial r^2} + \frac{1}{r}\frac{\partial[\text{H}_2\text{O}_2]}{\partial r} + \frac{\partial^2[\text{H}_2\text{O}_2]}{\partial x^2}\right). \tag{20}$$

The boundary conditions are:

$$\left.\frac{\partial[\text{H}_2\text{O}_2]}{\partial x}\right|_{x=0, r\leq R} = -\frac{\phi_c}{D_H}, \tag{21}$$

$$\left.\frac{\partial[\text{H}_2\text{O}_2]}{\partial x}\right|_{x=0, r\leq R} = 0, \tag{22}$$

$$\left.\frac{\partial[\text{H}_2\text{O}_2]}{\partial r}\right|_{r\to\infty} = 0, \tag{23}$$

$$\left.\frac{\partial[\text{H}_2\text{O}_2]}{\partial r}\right|_{r=0} = 0, \tag{24}$$

$$\left.\frac{\partial[\text{H}_2\text{O}_2]}{\partial x}\right|_{x\to\infty} = 0. \tag{25}$$

This is mathematically identical to the classical problem of a prescribed heat flux over a disk on an insulated semi-infinite solid [1], yielding the following solution in cylindrical coordinates $(r,\theta,x)$:

$$[H_2O_2](r,\theta,0)=\frac{1}{2\pi D_H}\int_0^{2\pi}\int_0^R \frac{\phi_c}{|r-r'|}r'dr'd\theta', \tag{26}$$

where

$$|r-r'|=\sqrt{r^2+r'^2-2\,r\,r'\cos(\theta-\theta')} \tag{27}$$

is the planar distance between the field point $(r,\theta,0)$ and the source point on the membrane plane, $(r',\theta',0)$. Setting $r=0$, the distance to any source point on the disk is simply the radial coordinate $r'$ of that point, so the double integral collapses to:

$$[H_2O_2]_{\max}=\frac{\phi_c}{2\pi D_H}\int_0^{2\pi}\int_0^R \frac{r'}{r'}dr'd\theta'=\frac{\phi_c}{2\pi D_H}2\pi R=\frac{\phi_c R}{D_H}. \tag{28}$$

#### 3. Image analysis and calculations to obtain the $H_2O_2$ landscapes in Figure 4

We took Figure 4A, left hand panel, of ref. [2], binarized it with an intensity threshold of 0.5, and obtained the centroid coordinates of all the spots (NOX2 clusters) with areas between 1 and 200 pixels. This yielded 811 clusters, in line with the reported average number 1 min post-stimulation. We then converted the centroid pixel coordinates to metric coordinates by scaling the former by the length of the scale bar in the image, which corresponds to 3 µm (private communication from Dr. Sophie Dupré-Crochet, Université Paris-Saclay, France). We computed the adhering area $(A)$ of the neutrophil as the area of the concave hull mesh of the set of selected clusters, setting $\alpha$= 0.83 as the lowest value that eliminated internal holes.

To compute the distribution of $H_2O_2$ at the surfaces of the neutrophil and of a receptor cell a given distance apart, we considered each cluster as a circle with 30 nm radius (half the mean reported diameter). Then, for each (x,y) point in the "landscape" we computed the distance to all cluster centroids, and used the concentration-distance relationships from the simulations shown in Figure 3B to sum the concentrations contributed to that point from all clusters. This operation applies the superposition principle, which is valid here due to the linearity of Model 4. Finally, we calibrated the obtained concentrations for the mean area-specific production rate to the reference maximal area-specific $H_2O_2$ production rate $\phi$ = 3.0 nmol dm$^{-2}$s$^{-1}$ value used for neutrophils in previous calculations. For this, we multiplied all the $H_2O_2$ concentrations by the factor

$$\gamma = \frac{\phi}{\dfrac{(n\ \text{cluster}) \times (16000\ \text{H}_2\text{O}_2\ \text{molecule cluster}^{-1}\text{s}^{-1})}{(A\ \text{dm}^2) \times 6.0 \times 10^{-14}\ \text{molecule nmol}^{-1}}}. \tag{29}$$

Image analysis and calculations described above, except the numerical simulations of the $H_2O_2$ distribution generated by a single NOX2 cluster, were performed in *Mathematica*™ v. 14.0.0.0 [3].

### 4. Supplementary figures

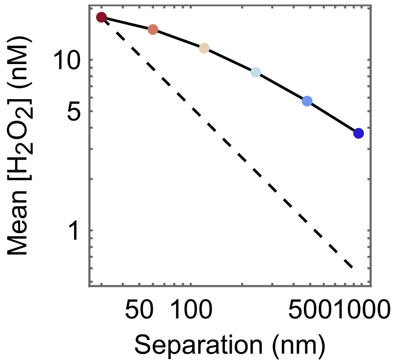

***Figure S 1. Mean $H_2O_2$ concentration over the surface of a 5 µm-radius receptor cell radially concentric with a 30 nm radius NOX2 cluster as function of the separation between source and receptor cells.*** *Calculations were based on Model 4. Note the log scales. The dashed line marks a reciprocal relationship. Support to Figure 3.*

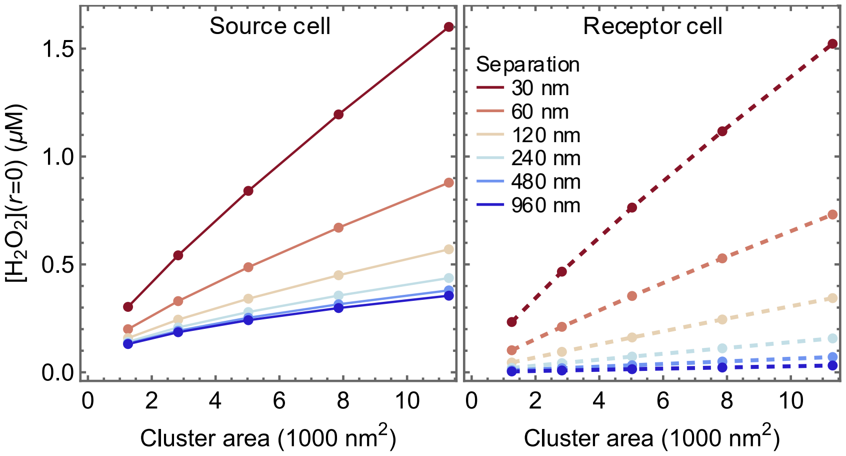

***Figure S 2. Maximal $H_2O_2$ concentrations near the surface of the source (solid lines, left) and receptor (dashed lines, right) cells near a fully active NOX2 cluster as function of the cluster area for various separations between the cells.*** *Calculations were based on Model 4. The lines are guides for the eye. Compare to Figure 3C inset.*

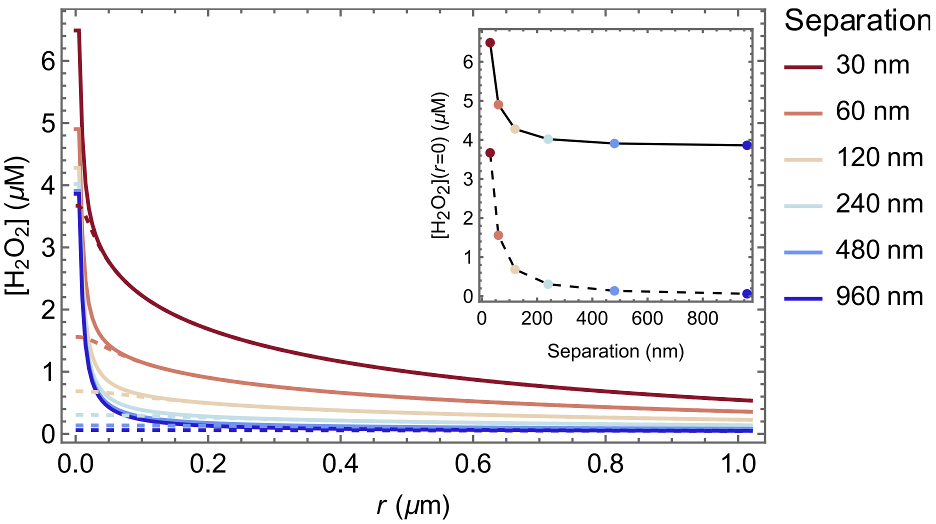

***Figure S 3. Distribution of $H_2O_2$ released from the neck of a NOX2-lined caveola to the ECS.*** *Distribution at the surface of the source (solid lines) and receptor (dashed) cells. Calculations were based on Model 4 with adaptations as described in the main text. The inset shows the dependence of the maximal concentration on the separation between the source and receptor cells.*

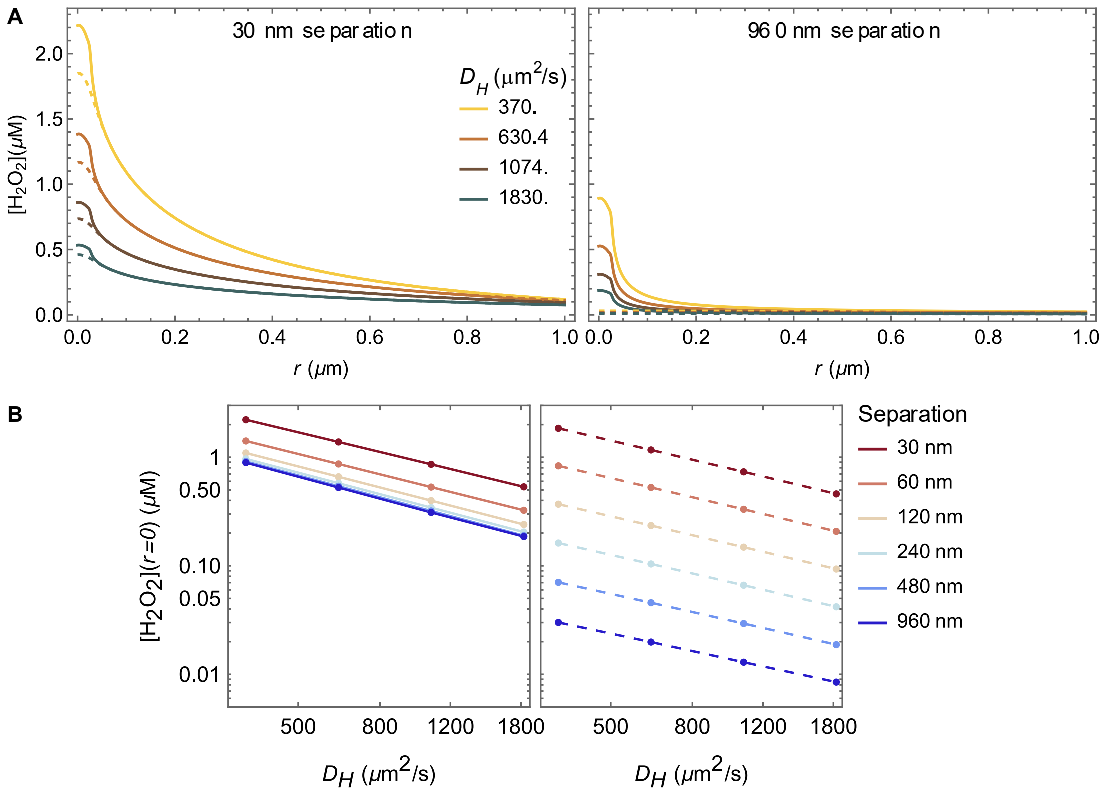

***Figure S 4. Effect of the diffusion coefficient on the distribution and maximal concentrations of $H_2O_2$ released from a 30 nm radius circular NOX2 cluster to the ECS. (A)*** *Distribution of $H_2O_2$ at the surface of the source (solid lines) and receptor (dashed) cells for 30 nm (left) or 960 nm (right) separation between the cells.* ***(B)*** *Log-log plots of the maximal $H_2O_2$ concentration at the surface of the source (left) and receptor (right) cells as function of the diffusion coefficient. The lines are guides for the eye. A $\phi_c$= 5.7×10$^6$ $H_2O_2$-equivalent molecules $s^{-1}\mu m^{-1}$ area-specific release rate by the clusters was assumed in all cases. Calculations were based on Model 4.*

***Supplementary Movie 1. Distribution of $H_2O_2$ (B) and of the Prdx (C-F), Trx (G,H), and glutathione (I,J) species over the first 5 minutes of a neutrophil active burst. (A)*** *Time course of $H_2O_2$ release from the source cell surface shown in orange in Figure 1B from the main text.* ***(B-J)*** *Concentration distributions over a vertical section cutting through the center of the geometry in Figure 1B. The horizontal white line marks the ECS, and the vertical lines mark the borders between the central cylindrical cells and the surrounding tissue. The color scale bars to the right of each plot represent the concentrations in μM units. Calculations were based on Model 6.*

[3] Wolfram Research Inc., Mathematica, Version 14, Wolfram Research, Inc., Champaign, IL, U.S.A., 2024.
