## Supplementary figures and images for "Concentration limits and localization of hydrogen peroxide in the extracellular space of solid tissues"

### Supplementary movie

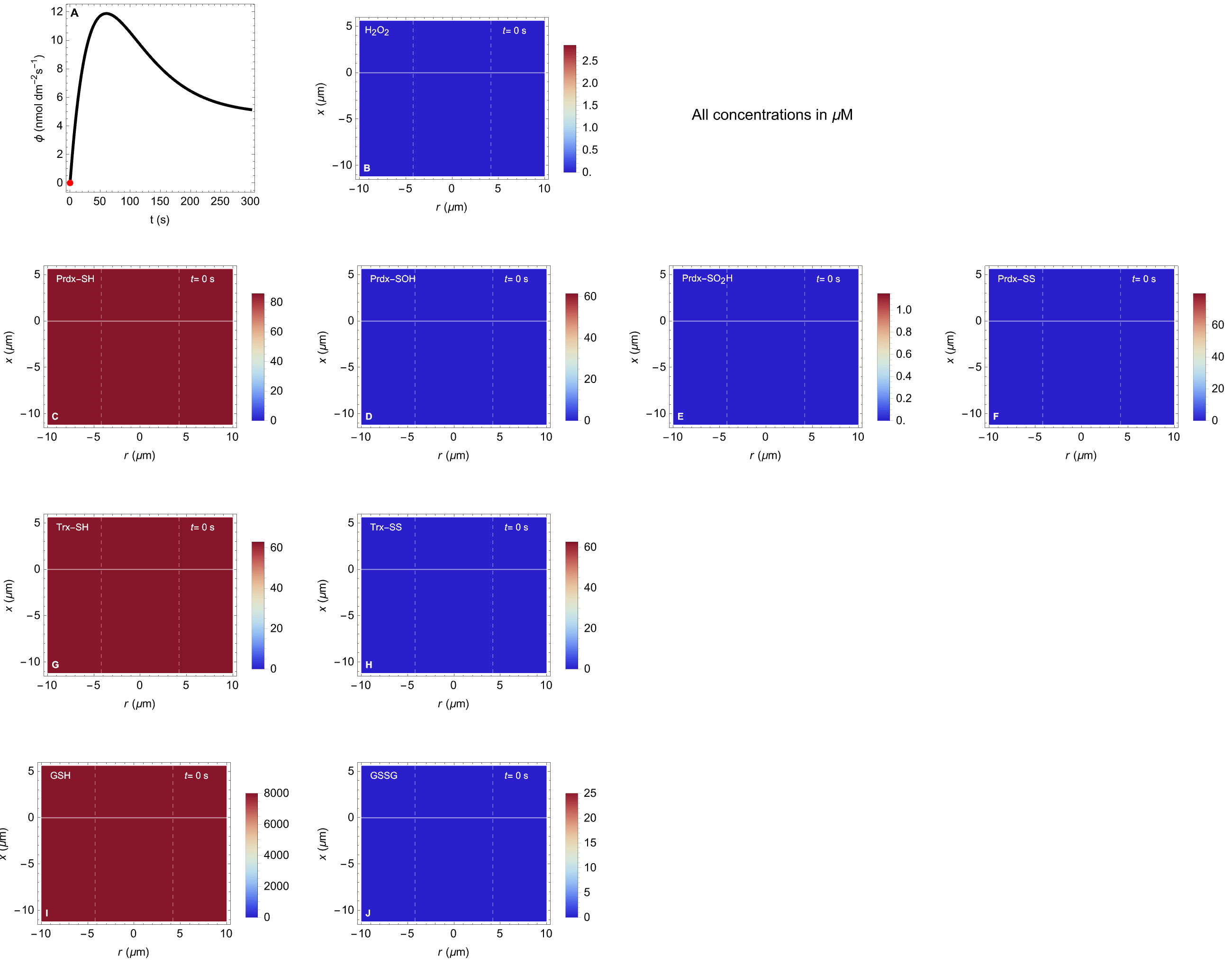
